# scConcept enables concept-level exploration of single-cell transcriptomic data

**DOI:** 10.64898/2026.04.21.719959

**Authors:** Hegang Chen, Yue Li

## Abstract

Interpreting high-dimensional single-cell transcriptomic data remains challenging, as existing methods rely on latent representations or prior knowledge that require extensive post hoc analysis to derive biologically meaningful insights. Topic models provide interpretable gene-level signals but often produce redundant and coarse-grained programs that are difficult to translate into coherent biological concepts. While recent foundation models and large language models (LLMs) show promise, they are not readily applicable to large-scale single-cell data or fail to provide structured, cell-level interpretations. Here we present scConcept, a framework that introduces concept-level representation by transforming gene-level topic representations into structured, human-interpretable biological concepts. By integrating neural topic modeling with LLMs, scConcept distills fragmented gene programs into semantically coherent concepts defined by a biological label, description, and gene set, and quantitatively maps them back to individual cells. Across 16 single-cell datasets, scConcept improves clustering performance by 27.1% and interpretability by 50.7% over state-of-the-art methods. These concept-level representations enable interpretable cell-state annotation and capture gene programs that generalize across datasets. In cancer applications, scConcept identifies clinically relevant programs associated with tumor progression and patient survival, and links them to candidate therapeutic targets. Together, scConcept establishes concept-level representation as a general and scalable abstraction for interpretable single-cell analysis.

## 1 Introduction

Gene expression profiling has been widely used to characterizing cells and tissues [1]. The rapid development of single-cell RNA sequencing (scRNA-seq) technologies has enabled researchers to resolve cellular composition, developmental stages, and disease-associated states at unprecedented resolution [2, 3]. A typical scRNA-seq dataset can be represented as a count matrix comprising approximately 20,000 genes and thousands to millions of cells. Analyzing and interpreting such datasets is a complex task that requires both bioinformatics expertise and domain-specific biological knowledge [4, 5]. A variety of computational tools have been developed for dimensionality reduction, cell clustering, differential expression analysis, and gene set analysis [6, 7]. In standard workflows, dimensionality reduction serves as a critical initial step, as the accuracy of capturing biologically relevant variation directly affects data visualization and downstream analyses, such as clustering for cell-type identification. Subsequent annotation is typically performed based on differential expression analysis. An alternative approach is interpretable dimensionality reduction, which explicitly models the contribution of individual genes to low-dimensional representations. By examining the genes contributing to each latent dimension, cell types can be directly interpreted without relying on clustering and differential expression analysis [8]. Compared with fully black-box approaches, such methods enhance interpretability and increase confidence in the biological relevance of the resulting representations.

Classical dimensionality reduction methods, such as principal component analysis (PCA), non-negative matrix factorization (NMF) and latent Dirichlet allocation (LDA), have been widely applied in single-cell analysis [9–11]. While the PCs lacks interpretability, NMF and LDA can produce interpretable basis factors each with explicit gene contributions per factor. However, these methods lack expressive capacity when applied to increasingly complex single-cell datasets. Another class of approaches is based on variational autoencoders (VAEs), such as scVI-LD [12] and neural topic models such as scETM [13], SPRUCE [14], d-scIGM [15], and scE^2^TM [16]. These models combine nonlinear encoders with linear decoders to achieve a balance between model expressiveness and interpretability. The latter further improve interpretability by learning the topic and gene embeddings on a shared manifold. Nonetheless, these methods face two major challenges. First, the learned topics are essentially collections of genes, and their interpretation often relies on downstream biological knowledge or gene set enrichment analysis. Specifically, the biological meaning of individual topics is not explicitly defined by the model and must be manually inspected post hoc. Almost in every dataset, some of the inferred topics tend to be enriched for a large number of heterogeneous biological pathways, which lack specificity and are often disregarded from downstream analysis. Second, even for the topics that do represent a meaningful pathway, many tend to be either too broad or redundant [17]. Although these topics capture low-level gene co-expression patterns, they do not form consistent and well-defined biological programs, thereby requiring substantial effort to interpret, validate, and distill biological insights from them. These limitations restrict the ability of topic models to directly support the understanding of cellular states and functions.

On the other hand, large language models (LLMs) have become promising tools for data analysis. LLMs are built upon the Transformer architecture, which employs self-attention mechanisms to model contextual information and capture long-range dependencies, enabling significant advances in tasks such as machine translation, topic modeling, and knowledge-based question answering [18–20]. These LLMs can help integrate information from large-scale corpora, including biomedical literature, and recent studies have further demonstrated their potential in bioinformatics applications, such as GenePT for constructing transferrable gene embedding [21] and gene set functional interpretation [22]. Building on these advances, LLMs provide important opportunities for analyzing and interpreting single-cell data. However, it is computationally expensive to directly train LLMs and perform inference using LLM on large-scale scRNA-seq data [23].

In this study, we present **scConcept**, an AI framework for automatic concept-level curation of single-cell transcriptomic data. scConcept builds upon neural topic models to learn gene-level topics from high-dimensional expression profiles and employs an LLM (e.g., GPT-5) as a domain expert to curate and turn these topics into human-interpretable biological concepts. Each concept is defined as a structured entity comprising a concise concept name, a natural language description, a biologically coherent gene set, and the set of source topics from which it is derived. By transforming fragmented and redundant gene-level signals to the concept level, scConcept substantially improves interpretability. Importantly, these concepts can be quantitatively mapped back to individual cells via their associated gene sets, enabling concept-level cell annotation. To systematically evaluate scConcept, we benchmark it against 10 state-of-the-art single-cell analysis methods across 16 scRNA-seq datasets, assessing both clustering performance and interpretability. scConcept improves clustering performance by 27.1% over the strongest baseline and interpretability by 50.7%. It captures biologically meaningful gene programs that are more coherent and less redundant, thereby enabling clearer characterization of cellular states. We further demonstrate the superiority of scConcept through a series of case studies. In a melanoma scRNA-seq dataset, scConcept generates concepts that closely correspond to annotated cell types and identifies tumor-associated programs. Concept-based *in-silico* perturbations simulate state transitions from malignant to normal cells. The identified concepts generalize to The Cancer Genome Atlas (TCGA) cohorts, where concept activity significantly correlates with patient survival, demonstrating clinical relevance. In hierarchical single-cell datasets, scConcept refines concepts into sub-concepts, enabling interpretable modeling of cellular hierarchies. Furthermore, scConcept accurately predicts cellular differentiation potential and discover transitions between developmental states through concept-based perturbations. Finally, we scale scConcept to a million-cell lung cancer atlas [24], where the identified concepts capture key components of the tumor microenvironment. These concepts generalize to independent TCGA cohorts, where their activity is significantly associated with clinical outcomes. Building on this, scConcept further enables concept-driven drug discovery by linking concept-derived gene programs to candidate therapeutic targets.

## 2 Results

scConcept integrates neural topic modeling with LLM to explore single-cell data in a more interpretable and effective manner. The overall workflow consists of three steps: (1) learning gene-level topics from cell expression data using a neural topic model; (2) distilling and synthesizing these topics into biological concepts using an LLM; (3) annotating cells with the derived concepts based on cell expression data. First, scConcept employs Embedding Clustering Regularization Topic Model (ECRTM) [25] to learn topics from high-dimensional single-cell expression profiles (Fig. 1A). By inheriting advantages from text modeling, ECRTM effectively handles sparse and high-dimensional data with ECR that mitigates topic redundancy, leading to more diverse topics. Each topic is subsequently represented as a gene set composed of the top 100 genes. Second, scConcept uses an LLM to distill these topics into biologically meaningful concepts (Fig. 1B). By filtering incoherent topics and consolidating redundant ones, the LLM derives a set of concepts, each defined as a structured entity comprising of a concise concept name, a natural language description, a coherent gene set, and the set of source topics that the concept was originated from. This process transforms fragmented topic representations into higher-level biological abstractions, improving interpretability for human experts. Finally, scConcept annotates cells by the concepts based on the expression of concept genes in each cell (Fig. 1B). Notably, this strategy obviates the need to predefine the number of clusters, as required in conventional clustering-based frameworks. Instead, the number of concepts is adaptively determined by the LLM, which captures diverse yet coherent biological signals from the learned topics. Therefore, scConcept combines the interpretable embedding capacity of neural topic models with the semantic abstraction capabilities of LLMs, thereby avoiding the high computational cost and context window limitations inherent to directly processing high-dimensional single-cell data with LLMs. Details are provided in the Section 4.1. In the following sections, we demonstrate the performance and real-world applications enabled by scConcept across a range of downstream tasks (Fig. 1C).

**Figure 1:**
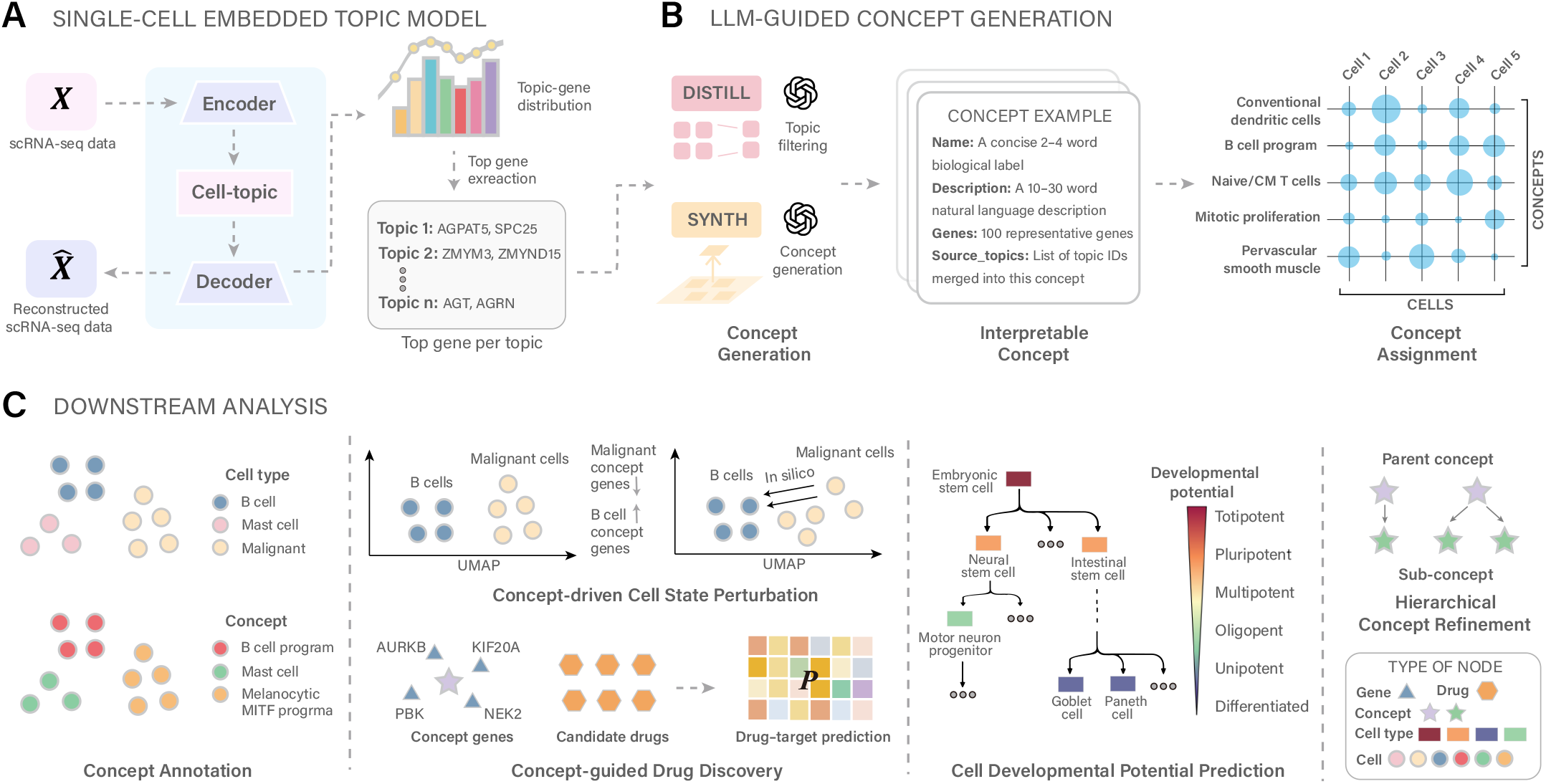
Overview of scConcept. (A) Topic extraction from single-cell data. A neural topic model is applied to scRNA-seq data to learn topic–gene distributions. For each topic, the top 100 genes with the highest weights are selected to characterize its underlying biological signal. (B) LLM-based concept generation. Topic gene sets are distilled using a large language model to filter incoherent topics and merge related ones into coherent biological concepts. Each concept is defined by a concise name, a natural language description, and a representative gene set. The resulting concepts are mapped back to individual cells based on gene expression, establishing a unified representation that links gene-level programs to cell-level states. (C) Concept-driven downstream analysis. Concept-level representations enable diverse analyses, including concept-based cell annotation, concept-guided cell state perturbation, concept-guided drug discovery, prediction of developmental potential, and hierarchical concept refinement.

### 2.1 scConcept enables accurate and interpretable cell-state representations

We evaluated scConcept against 10 leading single-cell methods across 16 scRNA-seq datasets spanning multiple species and sequencing technologies, with dataset sizes ranging from a few hundred to over 100,000 cells (Fig. 2A). We employed five quantitative metrics (Section 4.5) to evaluate clustering performance and interpretability, including Adjusted Rand Index (ARI), Normalized Mutual Information (NMI), Topic Coherence (TC), Topic Diversity (TD), and LLM-based Topic Coherence (TC-LLM). scConcept achieves the best overall performance (Fig. 2B), substantially outperforming the strongest competing method, scVI-LD, in terms of clustering (ARI and NMI; 27.1% performance gain; Wilcoxon signed-rank test, *p* = 6.1 *×* 10^−5^) and interpretability (50.7% performance gain; Wilcoxon signed-rank test *p* = 3.1 *×* 10^−5^).

**Figure 2:**
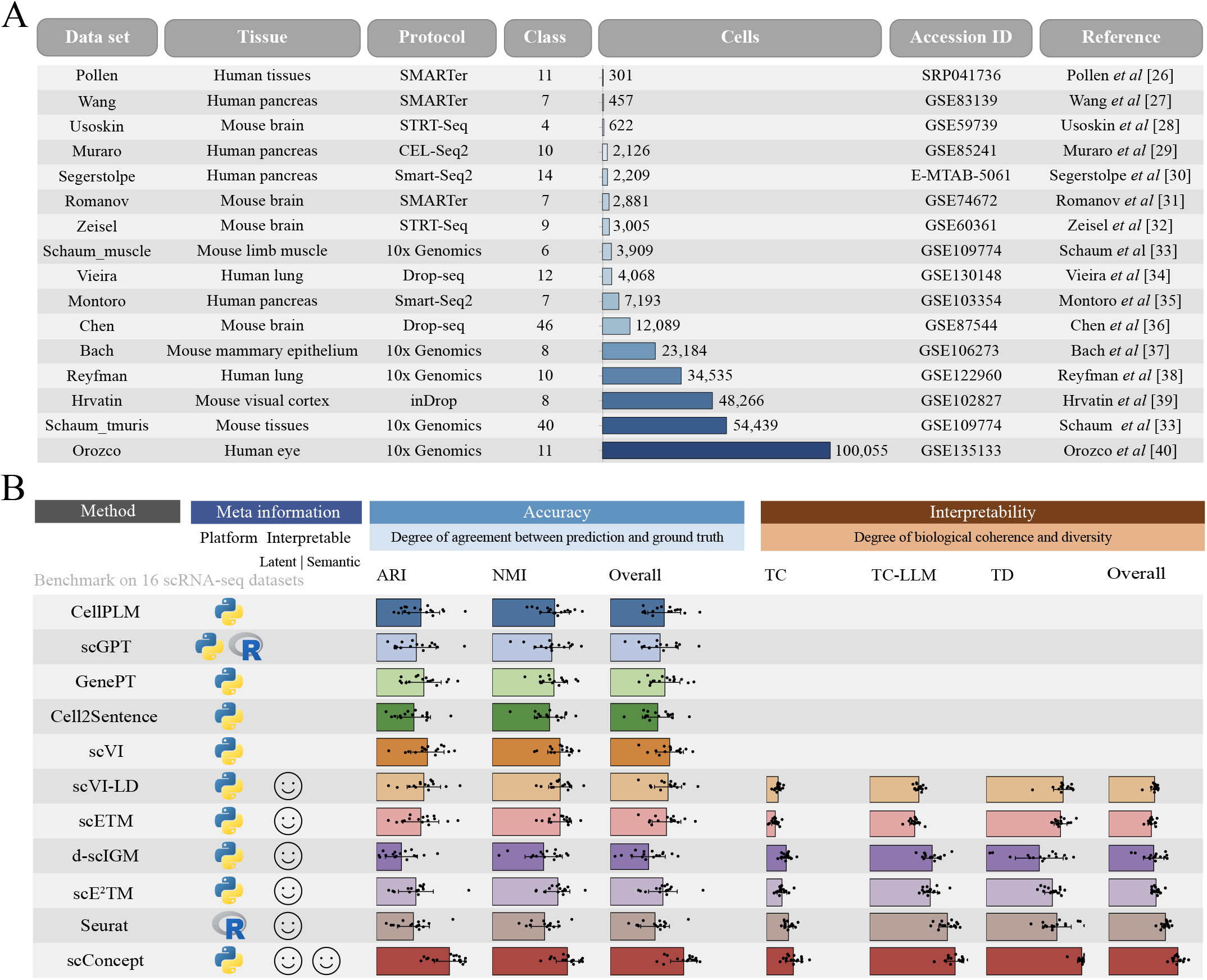
Clustering and interpretability benchmark. (A) Benchmark datasets. Summary of the 16 scRNA-seq datasets used in this study, including tissue source, sequencing protocol, number of annotated cell types, accession ID, and reference. Bar length is proportional to the number of cells in each dataset [26–40]. (B) Performance comparison across methods. Quantitative comparison of scConcept and baseline methods in terms of clustering accuracy and interpretability. Clustering accuracy is assessed by agreement with ground-truth annotations using ARI and NMI. Interpretability is evaluated using TC, TC-LLM, and TD, reflecting biological coherence, conceptual clarity, and diversity of the identified gene programs. Each point represents one dataset. The meta-information panel summarizes implementation platform (Python or R) and interpretability type, including *latent* interpretability (interpretation through latent variables) and *semantic* interpretability (human-readable concept-level descriptions).

Specifically, scConcept improves clustering performance by 23.5% compared to the strongest clustering baseline, scVI (Wilcoxon signed-rank test *p* = 3.1 *×* 10^−5^). For interpretability, it achieves an overall improvement of 21.2% across TC, TD, and TC-LLM metrics (Wilcoxon signed-rank test *p* = 3.1 *×* 10^−5^), outperforming the strongest interpretable baseline, Seurat. Methods such as d-scIGM, which explicitly incorporate prior biological knowledge, tend to exhibit strong topic coherence. However, these priors may overly constrain the optimization process, limiting the model’s ability to capture underlying data signals, resulting in the lowest clustering performance among the evaluated methods. Notably, scConcept substantially outperforms scE^2^TM, a method that integrates foundation model embeddings and ECR modules. This advantage stems from the ability of scConcept in transforming learned topics into coherent and non-redundant biological concepts and annotating individual cells with these concepts, providing a clearer and more interpretable characterization of cellular states.

Traditional clustering methods requires tunable resolution parameters (e.g., Louvain resolution). In contrast, scConcept directly assigns cells to concepts without requiring an explicit clustering step. The number of concepts is determined through LLM-based distillation, eliminating the need to manually specify cluster granularity. The results show that different baseline methods achieve their optimal clustering performance at different resolution settings (for example, scVI and Seurat at 0.1, and scETM at 0.3). Nevertheless, even across varied resolution settings, the best clustering performance achieved by baseline methods remains only comparable to that of scConcept, further supporting its advantage (Supplementary Fig. 1).

We additionally examined whether the improved interpretability of scConcept could be attributed to differences in the number of effective topics. Since scConcept performs topic filtering, it produce fewer concepts than the number of topics derived from the baseline models. To control for this factor, we evaluated all baseline methods under a reduced embedding dimensionality (set to 10) to match the effective number of concepts, where for topic-based models (e.g., scETM) the embedding dimensionality directly corresponds to the number of topics. As expected, reducing embedding dimensionality leads to a decrease in clustering performance for the baseline methods (Supplementary Fig. 2). Meanwhile, topic diversity and coherence increase due to the reduced number of topics. Nevertheless, scConcept maintains substantial advantages, achieving a 35.1% improvement over the strongest clustering baseline scVI (*p* = 6.1 *×* 10^−5^) and a 26.1% improvement over the strongest interpretability baseline scE^2^TM (*p* = 9.2 *×* 10^−5^). Overall, these results demonstrate that scConcept provides a unified framework for concept-level interpretation and representation of single-cell data in a concept-defined space, achieving strong performance in both clustering and interpretability.

### 2.2 scConcept captures transferable concepts

We hypothesized that the biological concepts identified by scConcept reflect underlying transcriptional programs and can generalize across datasets within the same biological context, particularly under cross-batch settings due to different experimental protocols or data sources. To evaluate whether scConcept captures transferable biological signals rather than dataset-specific patterns, we performed a series of cross-dataset transfer analyses. We considered six transfer tasks across related datasets: (1) human lung datasets (Reyfman→Vieira and Vieira→Reyfman), (2) human pancreas datasets (Wang→Muraro and Muraro→Wang), and (3) mouse brain datasets (Zeisel→Romanov and Romanov→Zeisel). For each task, genes were aligned between the reference and target datasets, and models were trained on the reference dataset only. The learned concepts were then directly applied to the target dataset without retraining.

We compared scConcept with three baseline methods that performed strongly in clustering, including scVI, scVI-LD, and scETM. Across all six transfer tasks, scConcept achieved the best overall performance (Supplementary Fig. 3). In particular, in the transfer from Muraro to Wang, scConcept attains substantially higher clustering accuracy (ARI: 0.876; NMI: 0.826) compared to scVI (ARI: 0.625; NMI: 0.717) and scETM (ARI: 0.635; NMI: 0.784). These results indicate that scConcept learns biologically meaningful and transferable concepts that capture shared transcriptional programs across related datasets.

### 2.3 scConcept reveals melanoma-associated concepts and enables perturbation and clinical relevance analysis

We applied scConcept to a melanoma scRNA-seq dataset [41] to identify concept-level gene programs and annotate cellular states. The dataset comprises 4,645 malignant, immune, or stromal cells isolated from 19 melanoma patients, including ten metastases to lymphoid tissues, eight distant-site metastases, and one primary acral melanoma. We first examined whether scConcept-derived concept annotations align with known cellular identities. UMAP visualization of the embedding space shows clear agreement between annotated cell types and concept assignments (Fig. 3A, B). For example, B cells are captured by a coherent *Mature B lymphocytes* concept, whereas fibroblast cells are distinctly separated from malignant cells and represented by an *Activated fibroblast matrix* concept. Notably, malignant cells are characterized by a specific *Melanocytic MITF program*. The LLM description of this concept is: *Pigmentation and melanocyte differentiation program with MITF, TYR, TYRP1, PMEL, and stress/secretory adaptations common in melanocytic tumor states*, therefore capturing a canonical transcriptional program associated with melanoma biology. Quantitative evaluation further confirms the superiority of scConcept in both clustering accuracy and interpretability (Fig. 3C). Together, these results demonstrate that scConcept captures biologically meaningful concepts that directly correspond to known cellular states for melanoma and can be reliably used to annotate individual cells.

**Figure 3:**
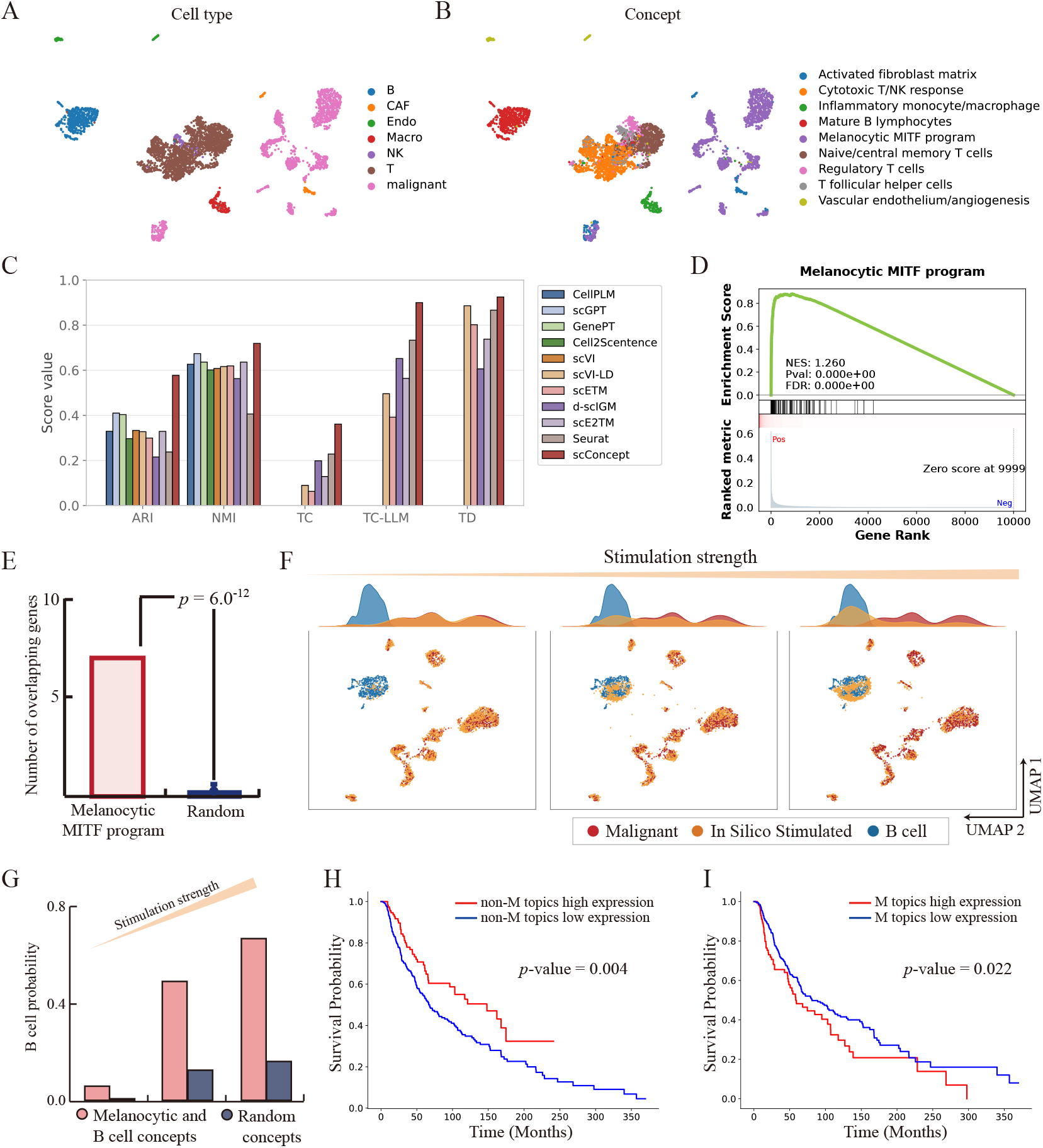
Concept-level analysis of melanoma reveals biologically meaningful programs, perturbation responses, and prognostic associations. (A) UMAP visualization of melanoma cells colored by annotated cell types. (B) UMAP visualization of melanoma cells colored by cell concepts. (C) Performance comparison across methods. Clustering accuracy and interpretability are compared across methods on the melanoma dataset using ARI, NMI, TC, TC-LLM, and TD. (D) Genes in the *Melanocytic MITF program* are significantly enriched in a ranked gene list derived from a logistic regression classifier, in which genes are ordered by their mean absolute SHAP values for malignant-cell prediction, indicating strong agreement between the concept and classifier-derived importance signals. (E) Enrichment against melanoma marker genes. Genes in the *Melanocytic MITF program* are significantly enriched for melanoma marker genes from the CellMarker database compared to randomly sampled gene sets of the same size. The expected overlap of random gene sets is estimated by averaging over 1,000 random samplings. Statistical significance is assessed using Fisher’s exact test. (F) Concept-driven cell-state perturbation. Malignant cells are perturbed by downregulating genes associated with the *Melanocytic MITF program* and upregulating genes associated with the *Mature B lymphocytes* concept. UMAP projections and density plots show progressive shifts in cell states with increasing perturbation strength. (G) Predicted B-cell probability under perturbation. The probability of perturbed malignant cells being classified as B cells increases with perturbation strength, as predicted by a logistic regression classifier trained on unperturbed malignant and B cells. (H) Survival analysis of non-malignant concepts. Kaplan–Meier survival curves for patients stratified by high versus low expression of concepts excluding the *Melanocytic MITF program* in the TCGA melanoma cohort. (I) Survival analysis of malignant-specific concepts. Kaplan–Meier survival curves for patients stratified by high versus low expression of the *Melanocytic MITF program* in the TCGA melanoma cohort.

We next focused on the tumor-associated concept identified by scConcept, the *Melanocytic MITF program*. A logistic regression classifier was trained to distinguish malignant from non-malignant cells, and genes were ranked according to their contributions to malignant-cell prediction using SHAP values [42]. Genes associated with the *Melanocytic MITF program* are significantly enriched among the top-ranked genes, indicating strong agreement between this concept and classifier-derived importance signals (Fig. 3D). Consistently, enrichment analysis against melanoma marker genes from the CellMarker [43] database shows significant overlap with genes in the *Melanocytic MITF program* (Fig. 3E), further supporting its biological validity. To investigate whether concept perturbations can drive cell-state transitions, we performed *in silico* perturbation by downregulating genes associated with the *Melanocytic MITF program* while upregulating genes associated with the *Mature B lymphocytes* concept (Section 4.8). As perturbation strength increases, the perturbed malignant cells progressively shift toward the B cell cluster in the embedding space (Fig. 3F). To quantify this transition, we trained a logistic regression classifier to predict cell states (malignant versus B cell). The fraction of perturbed malignant cells classified as B cells increases with perturbation strength (Fig. 3G), whereas perturbation of random concepts does not induce such transitions. These results suggest that the identified concepts capture key regulatory programs underlying cell-state identity and may represent potential drivers of phenotypic transitions.

To assess clinical relevance, we analyzed melanoma patient data from The Cancer Genome Atlas (TCGA) [44]. When transferring scConcept-derived concepts to TCGA gene expression data, we defined the *Melanocytic MITF program* as the malignant-associated concept, while all remaining concepts were considered non-malignant. Based on this definition, the expression levels of malignant- and non-malignant-associated concepts show significant associations with patient survival (Fig. 3H, I), with higher expression of the malignant-associated concept associated with reduced survival, whereas higher expression of non-malignant concepts is associated with improved survival. These findings indicate that scConcept identifies clinically relevant biological concepts that generalize across independent patient cohorts. Together, these results demonstrate that scConcept accurately identifies biologically meaningful concepts in melanoma and can be robustly extended to independent TCGA cohorts, providing an interpretable and reliable bridge between single-cell transcriptomics and cancer prognosis.

### 2.4 scConcept enables hierarchical modeling of cellular organization

With the increasing resolution of single-cell studies, cell types are increasingly defined at multiple levels [45, 46] and organized into hierarchical structures, enabling more fine-grained characterization of cellular states. To this end, we introduce a concept refinement procedure that extends scConcept to a hierarchical framework (Section 4.2). Briefly, for each concept, we assess whether the assigned cells exhibit substantial heterogeneity. Concepts showing pronounced heterogeneity are identified as divisible and further decomposed into sub-concepts using an LLM. Each subconcept is constrained to a subset of the parent concept’s gene set, enabling coherent biological interpretation across multiple levels of granularity.

We evaluated this hierarchical extension on two single-cell datasets with two-level annotations [47]. The human pancreas dataset contains 121,916 cells annotated into 9 coarse and 14 fine-grained cell types, while the human lung dataset comprises 318,426 cells with 44 coarse and 49 fine-grained annotations. At the coarse level, scConcept produces concept assignments that closely align with known cell-type identities (Fig. 4A, B). For example, pancreatic acinar cells are captured by a *Pancreatic acinar secretion* concept, and endothelial cells correspond to an *Endothelial angiogenesis* concept, demonstrating clear biological interpretability.

**Figure 4:**
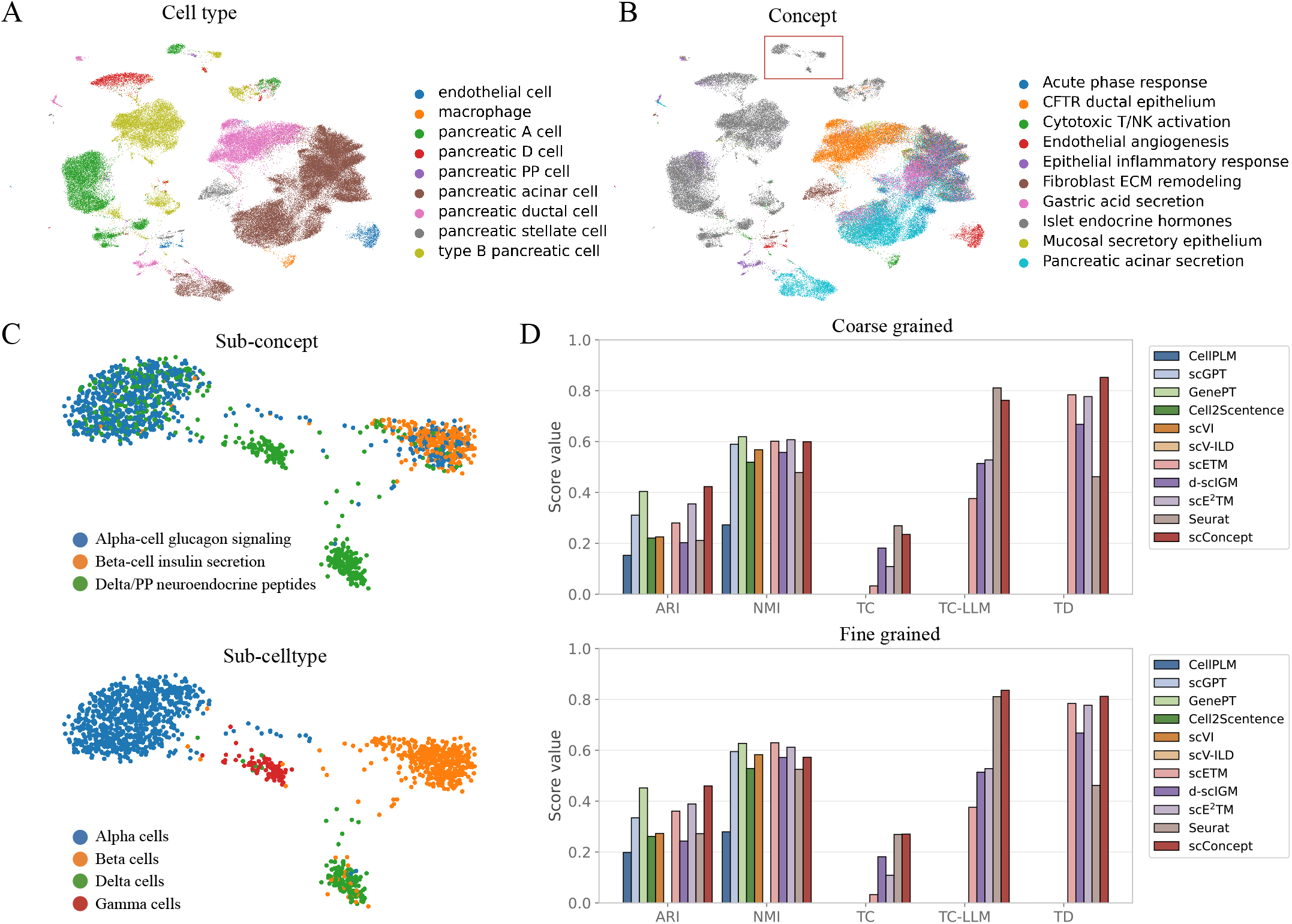
Hierarchical concept analysis reveals structured biological organization. (A) UMAP visualization of human pancreas cells colored by annotated cell types at the coarse grained level. (B) UMAP visualization of human pancreas cells colored by cell concepts. (C) Hierarchical refinement of concepts. Zoom-in visualization of the highlighted region in (B), showing refined sub-concepts (top) and corresponding fine grained cell-type annotations (bottom). (D) Performance comparison across granularities. Quantitative comparison of clustering accuracy and interpretability across methods at coarse grained and fine grained resolutions. Results are averaged over the pancreas and lung datasets.

At the fine-grained level, scConcept further refines the *Islet endocrine hormones* concept into more specific sub-concepts that correspond to distinct endocrine cell subtypes. These refined sub-concepts achieve improved alignment with ground-truth annotations (Fig. 4C). Notably, pancreatic PP cells, which account for only 0.28% of the dataset, are also accurately identified, highlighting the ability of scConcept to resolve rare cell populations. Quantitative evaluation confirms that scConcept achieves competitive clustering performance while consistently improving interpretability across both annotation levels (Fig. 4D). Together, these results demonstrate that scConcept effectively extends to hierarchical data, capturing multi-level biological organization and enabling coherent interpretation across different levels of cellular granularity.

### 2.5 scConcept predicts developmental potential and models cellular differentiation

Cells exhibit varying degrees of differentiation potential, defined as their capacity to generate specific cell types, spanning a continuum from totipotent and pluripotent states to multipotent, oligopotent, unipotent, and fully differentiated cells [48]. Although scRNA-seq technologies have advanced our understanding of cell fate, interpretable prediction of developmental potential remains a challenge. We applied scConcept to infer developmental potential at the single-cell level (Section 4.3). Specifically, gene programs associated with different stages of differentiation are extracted from the learned topics, giving rise to concepts that correspond to distinct developmental states. These concepts are then mapped to individual cells based on their expression profiles, enabling prediction of each cell’s developmental potential.

We collected six scRNA-seq datasets with annotated developmental potential from the study by Kang *et al* [49] (details in Supplementary Table 1). We compared scConcept with CytoTRACE2, a state-of-the-art supervised method for developmental potential prediction. The agreement between predicted and ground-truth annotations was quantified using Accuracy (Acc) and Kendall rank correlation (*τ*). Across all datasets, scConcept achieves performance comparable to Cyto-TRACE2 despite being unsupervised (Fig. 5A–C and Supplementary Fig. 4). In some datasets containing a single developmental potential label, scConcept accurately identifies the corresponding developmental stage, with Acc ≥ 0.98 (Supplementary Fig. 4). These results indicate that the concepts learned by scConcept capture biologically meaningful signals associated with different stages of differentiation, and that concept-level representations effectively characterize developmental potential at the single-cell level.

**Figure 5:**
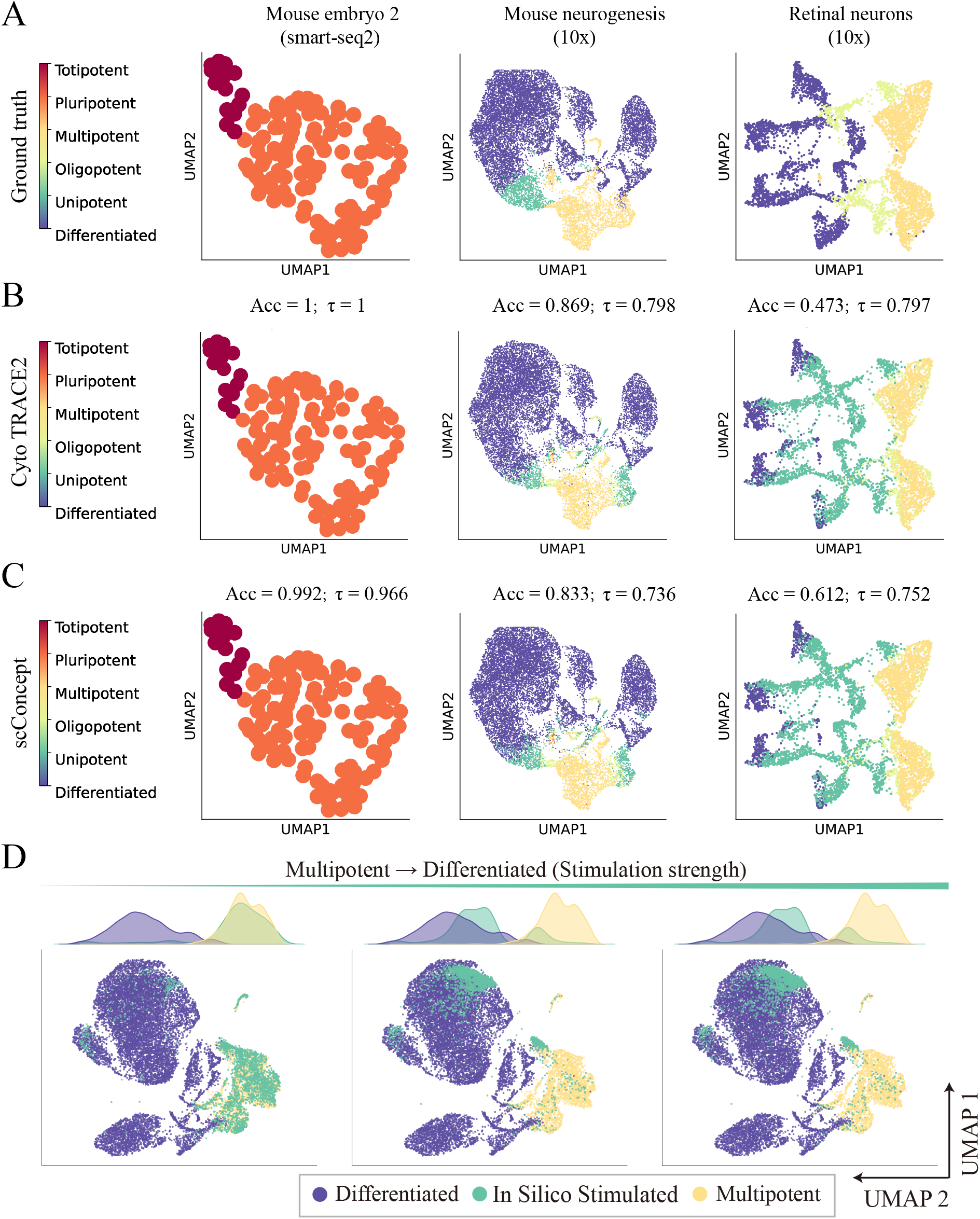
scConcept enables prediction of developmental potential and cell-state transitions. (A) Ground-truth developmental annotation. UMAP visualization of three representative datasets (mouse embryo, mouse neurogenesis, and retinal neurons) colored by annotated developmental potential, ranging from totipotent to differentiated states. (B) CytoTRACE2 predictions. UMAP visualization colored by developmental potential inferred by CytoTRACE2. Prediction performance is quantified by Accuracy (Acc) and Kendall rank correlation (*τ*), measuring agreement with ground-truth annotations. (C) scConcept predictions. UMAP visualization colored by developmental potential inferred by scConcept. (D) Concept-driven cell-state transitions. Cells are perturbed by downregulating genes associated with *multipotent* concept and upregulating genes associated with *differentiated* concept, simulating transitions from multipotent to differentiated states. UMAP projections and density plots show progressive shifts in cell states with increasing perturbation strength.

We further examined whether perturbing concepts associated with developmental stages can induce transitions between cellular states. Specifically, in a mouse neurogenesis dataset, we simulated perturbations in multipotent cells by downregulating genes associated with *multipotent* concepts (34 genes) while upregulating genes associated with *differentiated* concepts (44 genes). The full gene sets for each concept are provided in Supplementary Table 2. As perturbation strength increases, multipotent cells progressively shift toward differentiated states in the embedding space (Fig. 5D). To quantify this transition, we trained a logistic regression classifier to predict cell states (multipotent versus differentiated). The probability of perturbed multipotent cells being classified as differentiated increases accordingly (Supplementary Fig. 6). Conversely, when applying the opposite perturbation to differentiated cells, they progressively shift toward multipotent states in the embedding space, accompanied by an increased probability of being classified as multipotent (Supplementary Figs. 5 and 6). These results demonstrate that scConcept captures biologically meaningful concepts associated with different stages of differentiation, and that perturbing these concepts induces transitions between developmental states. This provides an interpretable framework for modeling and predicting developmental potential and facilitates a deeper understanding of cell potency.

### 2.6 scConcept links tumor microenvironment programs to clinical outcomes and drug discovery

Non-small cell lung cancer (NSCLC) exhibits substantial heterogeneity in its tumor microenvironment, which has been linked to disease progression and clinical outcomes. Large-scale single-cell atlases provide an opportunity to systematically characterize these microenvironmental programs and their clinical relevance. We applied scConcept to a large-scale single-cell transcriptomic atlas of NSCLC constructed by Salcher *et al*. [24], comprising 1.28 million single cells collected from 19 independent studies and 318 individuals. This dataset integrates diverse tumor and immune cell populations across multiple cohorts, providing a comprehensive resource for studying tumor microenvironment heterogeneity at scale. At the cellular level, scConcept produces concept assignments that are highly consistent with annotated cell types (Fig. 6A-C), demonstrating that concept-level representations remain robust even at this scale.

**Figure 6:**
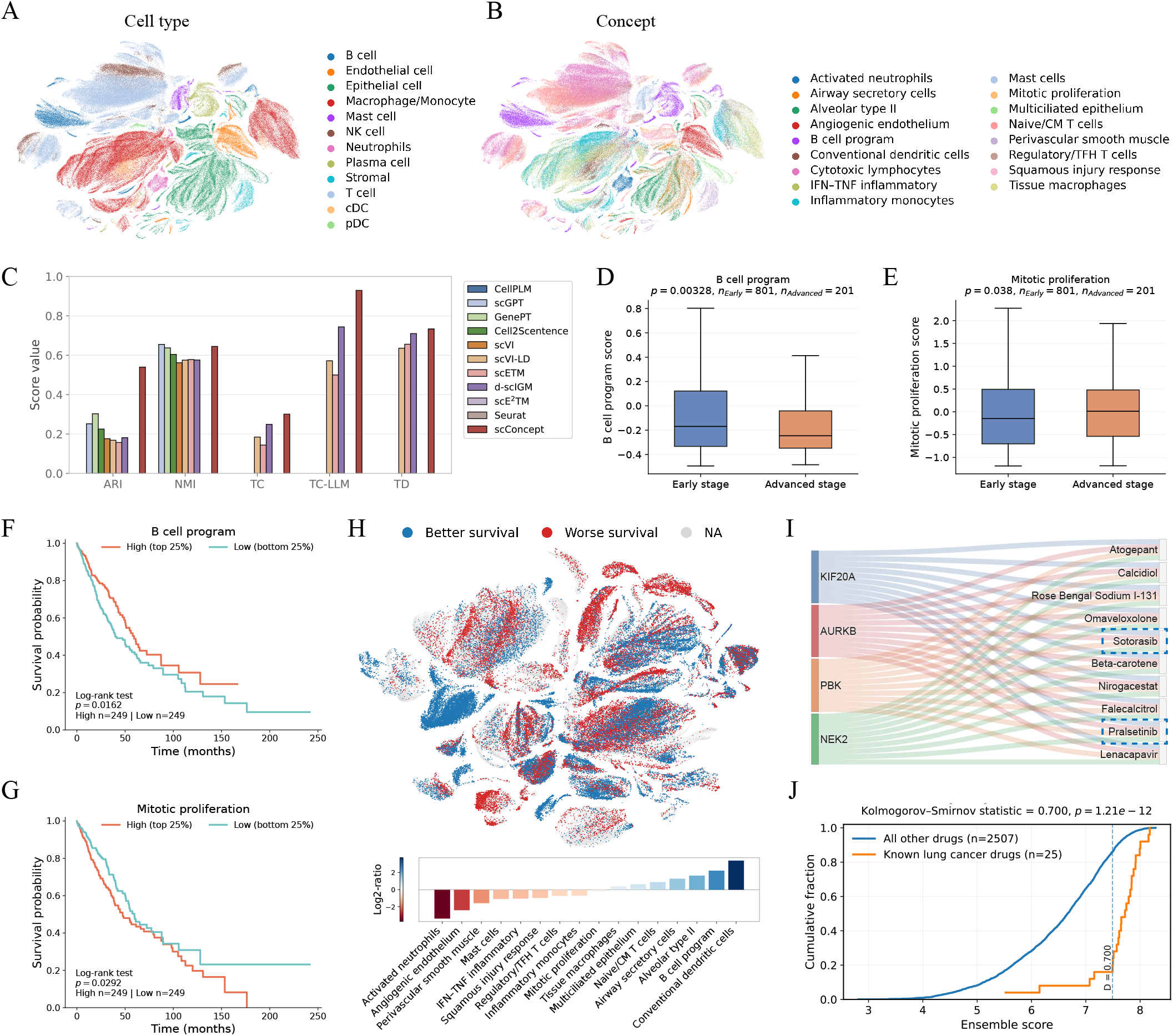
Concept-level analysis on a large-scale lung cancer atlas enables clinical and therapeutic insights. (A) UMAP visualization of the lung cancer atlas colored by annotated cell types. (B) UMAP visualization of the same cells colored by cell concepts. (C) Performance comparison across methods. Clustering accuracy and interpretability are compared across methods on the lung cancer atlas using ARI, NMI, TC, TC-LLM, and TD. (D) Stage-associated concept expression. The *B cell program* shows significantly higher expression in early-stage tumors than in advanced-stage tumors (Mann–Whitney U test). (E) Stage-associated concept expression. The *Mitotic proliferation* concept shows significantly higher expression in advanced-stage tumors than in early-stage tumors (Mann–Whitney U test). (F) Survival analysis of the *B cell program*. Kaplan–Meier survival curves show that patients with high expression of this concept exhibit improved survival compared to those with low expression. (G) Survival analysis of the *Mitotic proliferation* concept. Patients with high expression of this concept show reduced survival compared to those with low expression. (H) SCISSOR-based cellular association with survival. Integration of TCGA bulk RNA-seq phenotypes with single-cell data using SCISSOR. UMAP visualization highlights cells associated with better survival (scissor +, blue) or worse survival (scissor −, red). (I) Concept-guided drug–target associations. Sankey diagram showing predicted associations between target genes derived from the concept and the top 10 candidate drugs. Blue dashed boxes indicate drugs that are already FDA-approved for lung cancer treatment. (J) Comparison of drug ranking distributions. Cumulative distributions of predicted scores for known lung cancer drugs versus all candidate drugs. Differences between distributions are assessed using the Kolmogorov–Smirnov test.

We transferred scConcept-derived concepts to bulk RNA-seq data from 1,026 TCGA NSCLC patients, spanning UICC stages I–IV and including both lung adenocarcinoma (LUAD) and lung squamous cell carcinoma (LUSC). The *B cell program* and *Mitotic proliferation* concept exhibit strong associations with tumor stage (Fig. 6D, E). Specifically, the *B cell program* is enriched in early-stage tumors (stages I–II), whereas *mitotic proliferation* is elevated in advanced-stage tumors (stages III–IV). We next assessed the association between these two concepts and patient survival across 1,026 NSCLC patients. Survival analysis reveals that high expression of the *B cell program* is associated with improved patient outcomes, while elevated *mitotic proliferation* corresponds to poorer survival (Fig. 6F, G). Notably, these associations are statistically significant in LUAD but not in LUSC (Supplementary Figs. 7,8), which has been proposed in multiple previous studies [24, 50, 51]. To further characterize the association between scConcept-derived concepts and patient survival, we performed SCISSOR analysis [52] to integrate TCGA bulk phenotypes with single-cell data (Section 4.10). This analysis identifies cell populations whose transcriptional profiles are associated with survival. Immune-enriched cell populations, particularly B cells and dendritic cells, are positively associated with survival, whereas neutrophil populations show strong negative associations (Fig. 6H), consistent with established findings in tumor immunology. Importantly, these clinically relevant associations can be directly recovered from scConcept-derived annotations, without requiring manual cell-type labeling or additional deconvolution procedures, highlighting the ability of scConcept to provide a streamlined and interpretable framework for dissecting complex tumor microenvironmental signals.

Finally, we explored the potential of scConcept for interpretable drug discovery by focusing on a concept negatively associated with patient survival, the *Mitotic proliferation program*. We first examined the top 10 genes within this concept and found that 9 of them have previously been implicated in lung cancer (Supplementary Table 3), supporting the biological relevance of the identified program. We then mapped these genes to protein targets using the UniProt database [53] and queried drug–target interactions from the ChEMBL database [54]. Among these targets, four (KIF20A, AURKB, PBK, and NEK2) have known drug associations. To identify candidate therapeutics, we applied the state-of-the-art drug–target affinity prediction model PSICHIC [55] and screened 2,532 FDA-approved drugs against these targets (Section 4.12). The resulting ranking highlights several high-confidence candidates, among which two top-ranked drugs (e.g., Sotorasib and Pralsetinib) are already FDA-approved for lung cancer. providing strong support for the clinical relevance of the identified targets (Fig. 6I). We further evaluated the enrichment of known lung cancer drugs by comparing candidates derived from the Therapeutic Target Database [56] with all screened compounds. Known lung cancer drugs are significantly enriched among high-scoring candidates, as evidenced by a rightward shift in the cumulative distribution of predicted scores (Fig. 6J). This enrichment is statistically significant under the Kolmogorov–Smirnov test, indicating that scConcept-guided targets preferentially prioritize clinically relevant compounds. Together, scConcept enables the direct linking of tumor microenvironment-associated programs to patient survival and therapeutic targets, supporting concept-driven identification of clinically relevant drugs.

## 3 Discussion

Interpreting high-dimensional single-cell transcriptomic data in a biologically meaningful way remains a key challenge. While topic models provide a powerful framework for uncovering gene programs, their outputs are often fragmented, redundant, and require substantial domain expertise to interpret. In this work, we present scConcept, a framework that transforms gene-level topic representations into biologically meaningful concepts that are more readily interpretable by humans, and further maps these concepts back to individual cells to enable concept-level characterization of cellular states. Quantitative benchmarking demonstrates that scConcept achieves high performance in both clustering accuracy and interpretability. Notably, through a carefully designed integration of neural topic modeling and LLMs, scConcept avoids directly inputting high-dimensional single-cell expression data into LLMs, thereby circumventing the associated computational cost and context length limitations. This design establishes a practical and general framework for integrating LLM-based reasoning into single-cell analysis.

Beyond quantitative evaluation, scConcept demonstrates strong utility across diverse biological contexts. In single-cell melanoma data, scConcept identifies malignant-associated concepts and reveals their functional roles. Moreover, *in-silico* perturbation experiments show that modulating gene programs associated with specific concepts leads to consistent shifts in cellular states, thereby establishing a mechanistic link between gene programs and phenotypes. These concepts further generalize to independent melanoma patient cohorts from TCGA in terms of association with patient survival, demonstrating clinical relevance. Importantly, scConcept naturally extends to a range of downstream tasks. In hierarchical datasets, scConcept refines concepts into sub-concepts that align with multi-level cell-type annotations, enabling interpretable modeling of cellular hierarchies. In developmental systems, concept-based representations capture differentiation potential and reflect transitions between cellular states, providing insights into cellular potency. When scaled to a million-cell lung cancer atlas, scConcept uncovers clinically relevant associations among tumor stage, survival outcomes, and the tumor microenvironment, while also enabling concept-driven drug discovery. Together, these results demonstrate that scConcept provides a general framework for concept-level representation and interpretation of single-cell data, with strong performance and generalizability across diverse downstream tasks.

We also note some of scConcept ‘s limitations and areas for future development. First, the quality of concept distillation depends on the reliability of LLM outputs. Inaccuracies may arise in concept naming or gene composition. For example, in the human pancreas hierarchical dataset, macrophages cells are annotated with cytotoxic T activation (Fig. 4). Biologically, macrophages ingest and present antigens to T cells; activated cytotoxic T cells release cytokines (like IFN-*γ*) that further activate macrophages for enhanced killing. Although the two are key components of the immune system, the concept should have been more specific to macrophages than to cytotoxic T activation. One promising direction is to extend scConcept to a multi-agent framework, in which multiple LLM-based agents collaboratively refine concepts [5, 57, 58]. For instance, evaluation agents could assess the consistency between gene sets and concept descriptions, while filtering agents remove confounding genes, leading to more robust and reliable concept representations. Second, scConcept is currently designed for transcriptomic data and does not explicitly incorporate spatial context. Gene expression alone may be insufficient to capture the full complexity of cellular organization, particularly in tissues with strong spatial structure [59, 60]. Extending scConcept to spatial transcriptomics is therefore an important future direction. Integrating LLM-derived concept representations with spatially aware models could enable the incorporation of higher-level biological knowledge into spatial analysis, improving the identification of spatial domains and functional niches.

In summary, scConcept establishes an efficient AI framework to distill biologically meaningful concepts from single-cell transcriptomic data that are readily interpretable and can be harnessed to characterize cellular states for each cell. scConcept enables a wide range of downstream tasks from interpreting diverse cellular functions to discovering biological insights in relation to both basic biology and disease.

## 4 Methods

### 4.1 scConcept model

#### 4.1.1 Topic modeling

To extract topics from single-cell transcriptomic data, we adopt embedding clustering regularization topic model (ECRTM) as the backbone topic model. ECRTM is based on a VAE framework and incorporates an ECR component, which enables the learning of diverse and well-separated topic representations. Given an input cell expression vector **x**_*i*_ ∈ ℝ^*G*^, the encoder maps **x**_*i*_ to a latent Gaussian distribution parameterized by mean ***µ***_*i*_ and log-variance log ***σ***_*i*_:

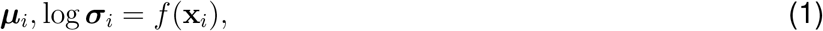

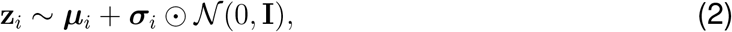

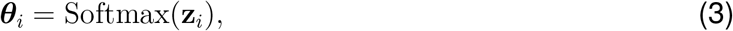

where ***θ***_*i*_ ∈ ℝ^*K*^ denotes the topic proportions of cell *i*.

The decoder reconstructs gene expression using the gene–topic matrix **B** ∈ ℝ^*G×K*^ as 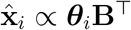. Each entry *b*_*mk*_ in **B** represents the association strength between gene *m* and topic *k*, and is parameterized based on the distance between gene embedding **g**_*m*_ and topic embedding **t**_*k*_:

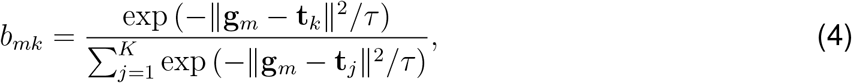

where *τ* is a temperature parameter that controls the sharpness of the gene–topic assignment.

To improve topic diversity, ECRTM incorporates ECR, which encourages gene embeddings to form well-separated clusters around topic embeddings. Further details of the ECR formulation and the ECRTM loss function are provided in Supplementary Notes 1 and 2.

#### 4.1.2 LLM-based concept generation

Given the learned gene–topic matrix **B** ∈ ℝ^*G×K*^, each topic is represented as a ranked list of genes according to their weights. For each topic *k*, a gene set 𝒢_*k*_ is constructed by selecting the top-*h* genes:

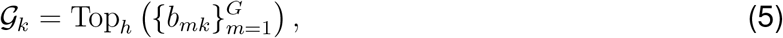

where *b*_*mk*_ denotes the weight of gene *m* in topic *k*, and *h* is set to 100 by default.

The collection of topic gene sets is denoted by 𝒢 = {𝒢_1_, …, 𝒢_*K*_}. These gene sets are provided as input to an LLM, which generates biologically coherent concepts by identifying shared gene programs across topics, merging redundant patterns, and filtering out incoherent or uninformative gene sets. This process can be expressed as 𝒞 = LLM(𝒢), where 𝒞 = {𝒞_1_, …, 𝒞_*M*_} denotes the resulting set of concepts. Each concept 𝒞_*j*_ is defined as a structured representation

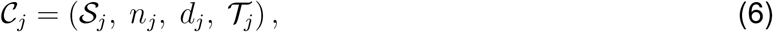

where 𝒮_*j*_ = {*m*_1_, …, *m*_*h*_} denotes the set of genes associated with the concept, *n*_*j*_ denotes the name of the concept, *d*_*j*_ provides a natural language description of the concept, and 𝒯_*j*_ denotes the set of source topics. In this study, we used GPT 5 as the LLM. Details of the prompt are provided in Supplementary Note 3.

#### 4.1.3 Cell–concept association

To quantify cell–concept associations, gene expression is first standardized across genes. Given the expression matrix **X** ∈ ℝ^*N ×G*^, where each row corresponds to a cell expression vector **x**_*i*_, the standardized expression 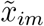 is computed as

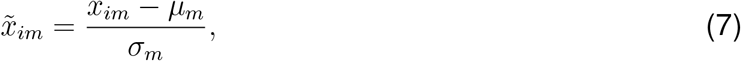

where *µ*_*m*_ and *σ*_*m*_ denote the mean and standard deviation of gene *m* across all cells. The cell–concept association score is defined as the average standardized expression over the concept gene set:

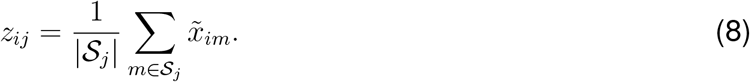

These scores form the cell–concept matrix **Z** ∈ ℝ^*N ×M*^, where *M* is the number of concepts. Each entry *z*_*ij*_ quantifies the strength of association between cell *i* and concept *j*. For discrete annotation, each cell is assigned to the concept with the highest score, *ĉ*_*i*_ = arg max_*j*_ *z*_*ij*_.

### 4.2 Hierarchical concept refinement

To capture biological structure at the higher granularity, scConcept extends the flat concept representation to a hierarchical formulation by recursively refining concepts into sub-concepts.

Given a concept 𝒞_*j*_ = (𝒮_*j*_, *n*_*j*_, *d*_*j*_, 𝒯_*j*_) and its associated cell–concept scores 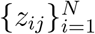, we first identify a subset of high-confidence cells associated with this concept using a score threshold:

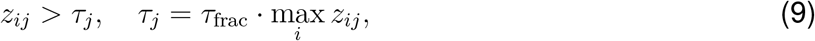

where *τ*_frac_ controls the fraction of the maximum score and is set to 0.5. Only concepts with at least a minimum number of associated cells are considered for refinement.

To determine whether a concept should be further split, we assess the heterogeneity of expression profiles within these high-confidence cells. Specifically, we restrict the expression matrix to the genes in the concept, forming a submatrix 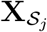. Let SSE_parent_ denote the total within-cluster sum of squared errors:

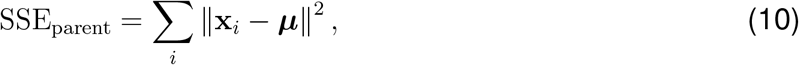

where ***µ*** is the mean expression vector of the selected cells. A candidate binary split is then obtained using KMeans clustering (*K* = 2), producing two subsets with corresponding errors SSE_1_ and SSE_2_. The impurity reduction is defined as

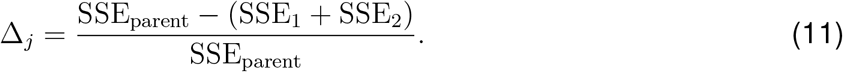

A concept is considered divisible only if (i) the number of high-confidence cells exceeds a minimum threshold, (ii) both child clusters satisfy a minimum leaf size, and (iii) the impurity reduction Δ_*j*_ exceeds a predefined threshold. In this study, *τ*_frac_ = 0.5, `min_cells` = 30, `min_leaf` = 20, and `min_impurity_reduction` = 0.1.

For concepts that satisfy these criteria, hierarchical refinement is performed using an LLM-based splitting operator, 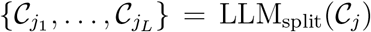, where each sub-concept 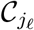 is defined in the same structured form as the parent concept. To ensure biological consistency, each sub-concept is constrained to select genes only from the parent gene set, thereby preventing the introduction of external genes during refinement. Further details of the prompt are provided in Supplementary Note 3.

After hierarchical refinement, concept assignment at deeper levels is performed in a structure-aware manner. For each cell *i*, the assignment at level *ℓ* is defined as

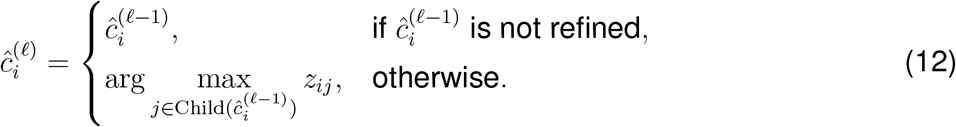

where *z*_*ij*_ denotes the assignment score between cell *i* and concept *j* at level *ℓ*. This hierarchical assignment ensures that fine-grained concept labels are selected within the local structure defined by their parent concept, preserving consistency across levels while enabling multi-resolution characterization of cellular heterogeneity.

### 4.3 Developmental concept extraction and cell annotation

To infer developmental potential at the single-cell level, scConcept extends the concept generation framework to identify gene programs associated with different stages of cellular potency. Starting from the learned topic gene sets, an LLM is prompted to distill gene programs corresponding to six predefined developmental potency categories, including totipotent, pluripotent, multipotent, oligopotent, unipotent, and differentiated states. This process is fully unsupervised and relies only on the provided topic gene lists without external marker databases or prior annotations. Because not all developmental stages are necessarily represented in a given dataset, some potency categories may not be used to annotate any topic gene set. Further details of the prompt are provided in Supplementary Note 3. Formally, the resulting set of potency concepts is denoted as

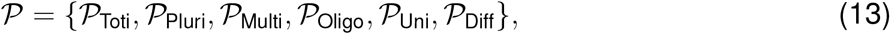

where each potency concept 𝒫_*ℓ*_ is defined as

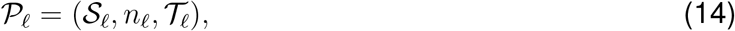

with *S*_*ℓ*_ denoting the gene set associated with potency level *ℓ, n*_*ℓ*_ denoting the predefined name of the potency category, and 𝒯_*ℓ*_ denoting the set of source topics supporting that concept.

To quantify the association between cells and developmental potency concepts, we compute a cell–concept score based on gene expression. Given the expression matrix **X** ∈ ℝ^*N ×G*^, the score between cell *i* and potency concept 𝒫_*ℓ*_ is defined as

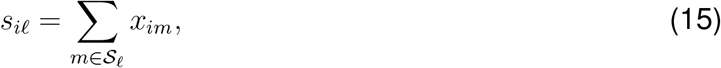

where *x*_*im*_ denotes the expression of gene *m* in cell *i*. Unlike the concept assignment procedure in Eq. 8, which averages over gene sets, this summation-based formulation accounts for both the magnitude of gene expression and the number of genes associated with each potency concept.

### 4.4 scRNA-seq datasets

Sixteen scRNA-seq datasets were used for benchmarking, including both human and mouse data. For all benchmarking datasets, we applied a unified preprocessing pipeline using the Scanpy package (v1.11.5). Specifically, we used `scanpy.pp.log1p()` to log-transform gene expression counts, followed by `scanpy.pp.highly_variable_genes()` to select the top 10,000 highly variable genes. Detailed information for all benchmarking datasets, including dataset name, tissue source, sample size, number of annotated cell types, sequencing protocol, and accession ID, is summarized in Fig. 2.

We further assembled multiple scRNA-seq datasets for downstream analyses spanning diverse biological contexts. For melanoma analysis, we used the dataset from Tirosh *et al*. [41] (GSE72056), as processed by Chen *et al*. [15], resulting in 4,092 cells across seven cell types with 10,000 highly variable genes. For hierarchical analysis, we used two scRNA-seq datasets with coarse- and fine-grained cell-type annotations from Xu *et al*. [47]. The pancreas dataset contained 121,916 cells with 9 coarse-grained and 14 fine-grained cell types, whereas the lung dataset contained 318,426 cells with 44 coarse-grained and 49 fine-grained cell types. For both datasets, we applied the same preprocessing procedure to log-transform gene expression and to select the top-10,000 highly variable genes. The original data are available at https://www.celltypist.org/organs. For developmental potential analysis, we collected six scRNA-seq datasets with annotated developmental stages from Kang *et al*. [49]. All datasets were processed using the same preprocessing procedure, including log-transformation and selection of the top-10,000 highly variable genes. Detailed dataset descriptions are provided in Supplementary Table 1. The original data are available at https://cytotrace2.stanford.edu. For large-scale analysis, we used the non-small cell lung cancer atlas from Salcher *et al*. [24], which integrates scRNA-seq data from 19 independent studies, comprising 1,283,972 cells from 318 individuals. We used the preprocessed data provided by the authors, including 6,000 highly variable genes. The original data are available at https://zenodo.org/records/7227571.

### 4.5 Benchmark metrics

For evaluating cell clustering, we used standard metrics, namely the Normalized Mutual Information (NMI) and the Adjusted Rand Index (ARI). All embedding maps were visualized using UMAP, and cells were clustered using Louvain’s algorithm [61]. For developmental potential prediction, we evaluated performance using Accuracy (Acc) and Kendall rank correlation coefficient (*τ*).

To assess interpretability, we used Topic Coherence (TC), Topic Diversity (TD), and a Large Language Model-based Topic Coherence metric (TC-LLM) as detailed below.

#### TC

Topic coherence measures the semantic consistency of genes within each topic by evaluating their co-occurrence patterns in biological pathways. For the calculation of TC, we used as the external corpus all known gene pathways for human and mouse, namely `msigdb.v2024.1.Hs.symbols` and `msigdb.v2024.1.Mm.symbols`, respectively, downloaded from the GSEA database [62]. Formally,

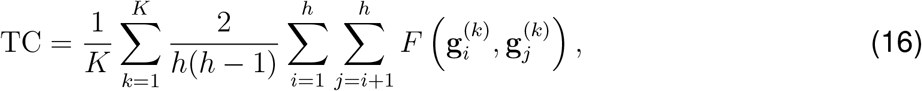

where 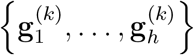 denotes the top-*h* genes of the *k*-th topic or concept. TC ranges from 0 to 1, with higher values indicating stronger coherence.

#### TD

Topic diversity quantifies the distinctness of biological signals captured by different topics. It is defined as

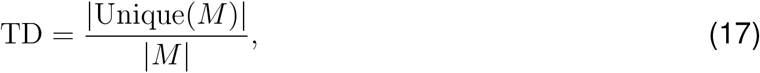

where 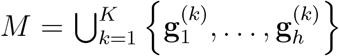 is the union of top-*h* genes across all topics or concepts. TD ranges from 0 to 1, with higher values indicating greater diversity and lower redundancy.

#### TC-LLM

To further assess interpretability, we introduce TC-LLM, a large language model-based coherence metric. For each topic or concept, we provided the associated gene set to an LLM and asked it to assign a coherence score on a five-level scale (1: not interpretable; 5: highly coherent, representing a clear biological program, pathway, or cell identity). The detailed prompt is provided in Supplementary Note 3. The resulting scores were linearly normalized to the range [0, 1].

For TC, TD, and TC-LLM, we followed previous work [16] and used the top 10 genes for each topic. For scConcept, these correspond to the first 10 genes listed in each LLM-generated concept gene set.

### 4.6 Baseline methods

Single-cell foundation models have received increasing attention in recent years. We compared scConcept with four representative methods, including CellPLM [63], scGPT [64], GenePT [21], and Cell2Sentence [23]. For GenePT, cell-level embeddings were constructed by computing a weighted average of gene embeddings based on gene expression levels.

For interpretable single-cell embedding models, we considered scVI-LD [12], scETM [13], d-scIGM [15], and scE^2^TM [16] as baselines. scVI-LD method provides interpretability by decomposing the decoder to establish relationships between cell representations and gene weights, where the cell latent coordinates are treated as topics. scETM model introduces interpretable linear decoders to learn biologically meaningful topic and gene embeddings. d-scIGM model incorporates hierarchical prior knowledge and deeper neural architectures to capture complex dependencies between genes and topics; to ensure fair comparison, we used the version without prior knowledge as described in the original study. scE^2^TM model introduces ECR, which encourages each topic to represent a distinct group of genes and enables the learning of diverse and biologically informative topics. In addition, we included scVI [65] as a baseline, a classical deep generative model for single-cell data that has demonstrated strong performance in previous benchmarks [66]. We also considered a cluster-based differential expression pipeline implemented with Seurat [11].

### 4.7 Implementation details

scConcept is implemented in Python and built upon the PyTorch (v2.9.0) framework. We adopt ECRTM as the backbone model. During training, the number of topics is fixed to *K* = 50. The hidden dimension of each layer is set to 200, and the `tanh()` function is used as the activation. The model is optimized using the RMSprop optimizer for 500 iterations. For datasets with more than 10,000 cells, a batch size of 2,048 is used; otherwise, the batch size is set to 512. The regularization weight *λ* is fixed to 100. The entropic regularization parameter *ϵ* in the optimal transport formulation is set to 20, following Wu *et al*. [25]. All LLM-based components in this study were implemented using GPT-5. To improve reproducibility, we fixed the random seed to 1 for all LLM queries. All UMAP visualizations in this study were generated using Scanpy (v1.11.5). Specifically, principal component analysis (PCA) was first applied to the input gene expression data, followed by neighborhood graph construction and UMAP embedding using default parameters unless otherwise specified.

To ensure fair comparison across methods, we adopted a unified clustering pipeline for all embedding-based approaches. For single-cell foundation models, including CellPLM, scGPT, GenePT, and Cell2Sentence, the learned embeddings were first reduced to 50 dimensions using `scanpy.tl.pca`. For all other embedding-based methods, including scVI, scVI-LD, scETM, d-scIGM, and scE^2^TM, the latent embedding dimension was directly set to 50. For the embeddings obtained from these methods, clustering was performed by constructing a neighborhood graph using `scanpy.pp.neighbors`, followed by Louvain clustering using `scanpy.tl.louvain`. For the Seurat-based pipeline, clustering was performed using the standard workflow, including `RunPCA` (with 50 components), `FindNeighbors`, and `FindClusters` (Louvain algorithm). Differentially expressed genes for each cluster were identified using `FindAllMarkers` (Wilcoxon rank-sum test), and the resulting gene sets were treated as gene programs for interpretability comparison. All baseline models were implemented using publicly available code repositories. Hyperparameters not explicitly specified were set to default values as reported in the original publications or corresponding codebases. Most experiments were conducted on a system equipped with an Nvidia RTX 2080 Ti GPU and 378 GB RAM. For large-scale datasets, including the human lung hierarchical dataset and the non-small cell lung cancer atlas, computations were performed on an Nvidia A100 GPU (40 GB memory).

### 4.8 Concept-based *in silico* perturbation experiments

We performed concept-based *in silico* perturbation experiments by directly modifying gene expression profiles according to concept-associated gene sets. For concept upregulation, the expression values of genes belonging to a given concept were amplified in each cell by multiplying their original expression levels by factors of 8, 32, and 64, simulating increasing strengths of concept activation. For concept downregulation, the expression values of the same genes were attenuated by scaling them to 1*/*8, 1*/*32, and 1*/*64 of their original values, representing progressive suppression of the corresponding biological program. To evaluate the specificity of the perturbations, we performed negative control experiments by randomly selecting an equal number of concepts and applying the same upregulation and downregulation procedures.

### 4.9 Categorization of perturbed cells

To evaluate the effect of perturbations on cellular states, we first divided the unperturbed dataset into training and testing sets in a 7:3 ratio. A logistic regression classifier, implemented using `scikit-learn`, was trained on the unperturbed training set to predict the treatment or condition label of each cell. After training, we identified the subset of perturbed cells corresponding to the original testing set and applied the trained classifier to these cells for prediction. The reported results represent the average performance over 10 independent runs.

### 4.10 SCISSOR analysis

We used SCISSOR [52] to associate phenotypic data from bulk RNA-seq experiments with our single-cell data. Following the analysis workflow described by Salcher *et al*. [24], we applied SCISSOR to primary tumor cells of each patient individually, using overall survival (Cox regression) as the dependent variable. Default parameters were used for all SCISSOR analyses.

A total of 21 out of 176 samples with low overall cell counts failed during the Seurat preprocessing step of SCISSOR and were excluded from subsequent analyses. For each cell type, we computed the log_2_ ratio of scissor + and scissor − cells, defined as the mean fraction of scissor + cells divided by the mean fraction of scissor − cells. Statistical significance was assessed by comparing the fractions of scissor + and scissor − cells using a paired Wilcoxon test implemented in the `scipy` package. *P*-values were adjusted using the Benjamini–Hochberg procedure and considered significant at a false discovery rate (FDR) < 0.01.

### 4.11 Survival analysis of cancer patients from TCGA

We transferred scConcept-derived concepts to melanoma and lung cancer patient cohorts and performed survival analysis. For melanoma, we extracted clinical and bulk gene expression data from the TCGA database (cancer study id: skcm_tcga) using the `cgdsr` R package. Concepts identified from single-cell data were projected onto bulk RNA-seq data by computing patient-level concept scores based on gene expression profiles, where each concept was represented using the first 10 genes in the corresponding LLM-generated gene set. We defined the *Melanocytic MITF program* as a malignant-associated concept, while all other concepts were considered non-malignant. Patients were stratified into two groups based on concept scores, corresponding to low (bottom 25%) and high (top 25%) expression levels. Kaplan–Meier survival curves were estimated using the `survival` R package, and differences between groups were evaluated accordingly. For lung cancer, we used survival data from Liu *et al*. [67]. The same concept transfer and survival analysis procedure was applied.

### 4.12 Drug–target affinity prediction

To identify candidate therapeutics associated with the *Mitotic proliferation program*, we performed drug–target affinity prediction based on four genes (KIF20A, AURKB, PBK, and NEK2). Specifically, genes within the concept were first mapped to their corresponding protein sequences using the UniProt database [53]. The resulting protein sequences were represented in FASTA format and used as inputs for downstream prediction. Candidate drugs and reference compounds were obtained from the Therapeutic Target Database [56]. For each drug, the chemical structure was represented using its SMILES string retrieved from the database. Drug–target interactions were then evaluated using PSICHIC [55], a deep learning framework that predicts protein–ligand binding affinity directly from sequence information. PSICHIC takes as input the protein amino acid sequence and the molecular SMILES representation of the ligand, and outputs a quantitative affinity score reflecting interaction strength. For each target, affinity scores were computed across all candidate drugs. To obtain a unified drug ranking, the predicted scores were averaged across the four targets, yielding a final affinity score for each drug.

### 4.13 Code availability

The scConcept model and all the code necessary for reproducing our results is publicly available via GitHub at https://github.com/li-lab-mcgill/scConcept. To ensure reproducible results, all analyses are performed using the same fixed random seed. An archived version will be deposited in the Zenodo database upon acceptance.

## Supporting information

Includes supplementary notes, supplementary figures, and supplementary tables.

## 5 Acknowledgments

Y.L. is supported by Canada Research Chair (Tier 2) in Machine Learning for Genomics and Healthcare (CRC-2021-00547) and CIHR Project Grant (PJT-540722).

