## Supplementary material for "scConcept enables concept-level exploration of single-cell transcriptomic data": Includes supplementary notes, supplementary figures, and supplementary tables.

### Supplementary materials

#### Contents

**1 Supplementary Notes** **2**

**2 Supplementary Figures** **10**

**3 Supplementary Tables** **16**

### 1 Supplementary Notes

#### Supplementary Note 1: Embedding clustering regularization

ECRTM [1] incorporates an embedding clustering regularization (ECR) to model gene–topic relationships. In the context of single-cell transcriptomic data, each topic embedding $\mathbf{t}_k$ acts as a centroid of a gene cluster, while gene embeddings $\mathbf{g}_m$ represent data points. The soft assignment of genes to topics is modeled using an entropy-regularized optimal transport (OT) framework. Specifically, $K$ topic embeddings serve as cluster centroids and $V$ gene embeddings correspond to data points. Let $n_k$ denote the number of genes assigned to topic $k$, and $s_k = n_k/V$ denote the corresponding cluster proportion. The vector $\mathbf{s} = (s_1, \ldots, s_K)^\top \in \Delta_K$ summarizes the cluster proportions. A uniform prior over clusters is assumed, i.e., $s_k = 1/K$, which avoids rivial669 solutions with empty clusters [1, 2]. Two discrete measures over gene and topic embeddings are defined as $\gamma = \sum_{m=1}^{V} \frac{1}{V}\delta_{\mathbf{g}_m}$ and $\phi = \sum_{k=1}^{K} s_k \delta_{\mathbf{t}_k}$, where $\delta_z$ denotes the Dirac measure at $z$. The entropy-regularized OT problem between $\gamma$ and $\phi$ is formulated as

$$\underset{\boldsymbol{\pi}\in\mathbb{R}_+^{V\times K}}{\arg\min}\mathcal{L}_{\mathrm{OT}_\varepsilon}(\gamma,\phi) = \sum_{m=1}^{V}\sum_{k=1}^{K} \mathrm{D}\left(\mathbf{g}_m, \mathbf{t}_k\right)\pi_{mk} + \varepsilon \sum_{m=1}^{V}\sum_{k=1}^{K} \pi_{mk}\left(\log \pi_{mk} - 1\right)$$

$$\text{s.t.} \quad \boldsymbol{\pi}\mathbf{1}_K = \frac{1}{V}\mathbf{1}_V, \quad \boldsymbol{\pi}^\top \mathbf{1}_V = \frac{1}{K}\mathbf{1}_K, \tag{1}$$

where $\varepsilon$ controls the strength of entropy regularization. The transport cost is defined as the squared Euclidean distance,

$$\mathrm{D}(\mathbf{g}_m, \mathbf{t}_k) = C_{mk} = \|\mathbf{g}_m - \mathbf{t}_k\|^2, \tag{2}$$

and $\mathbf{C} \in \mathbb{R}^{V\times K}$ denotes the corresponding cost matrix. The constraints in Eq. (1) ensure that each gene embedding has total mass $1/V$ and each topic embedding has total mass $1/K$. The resulting transport weights $\pi^*_{mk}$ define the soft assignment of gene $\mathbf{g}_m$ to topic $\mathbf{t}_k$. Besides, $\pi_{mk}$ denotes the transport weight from $\mathbf{g}_m$ to $\mathbf{t}_k$ and $\boldsymbol{\pi} \in \mathbb{R}_+^{V\times K}$ is the transport plan that includes the transport weight of each gene embedding to fulfill the weight of each topic embedding. To meet these constraints, ECR models clustering soft assignments with the optimal transport plan $\boldsymbol{\pi}^*_\varepsilon$, i.e., the soft-assignment of $\mathbf{g}_m$ to $\mathbf{t}_k$ is the transport weight between them, $\pi^*_{\varepsilon,mk}$. The formula is defined as follows:

$$\mathcal{L}_{\mathrm{ECR}} = \sum_{m=1}^{V}\sum_{k=1}^{K} \|\mathbf{g}_m - \mathbf{t}_k\|^2 \pi^*_{mk}. \tag{3}$$

where

$$\pi^*_\varepsilon \leftarrow \mathrm{sinkhorn}(\gamma, \phi, \varepsilon) \approx \underset{\boldsymbol{\pi}\in\mathbb{R}_+^{V\times K}}{\arg\min}\mathcal{L}_{\mathrm{OT}_\varepsilon}(\gamma,\phi) \tag{4}$$

This formulation encourages gene embeddings to form well-separated clusters around topic embeddings, thereby improving topic diversity.

#### Supplementary Note 2: ECRTM loss function

The training loss of ECRTM follows the standard variational autoencoder (VAE) framework. The VAE loss is defined as

$$\mathcal{L}_{\text{VAE}} = \frac{1}{n}\sum_{i=1}^{n} -\mathbf{x}_i^\top \log\left(\text{Softmax}\left(\boldsymbol{\theta}_i \mathbf{B}^\top\right)\right) + \text{KL}\left[q\left(\boldsymbol{\theta}_i \mid \mathbf{x}_i\right) \| p\left(\boldsymbol{\theta}_i\right)\right], \quad (5)$$

where the first term corresponds to the reconstruction error, and the second term is the Kullback–Leibler (KL) divergence between the variational posterior and the prior distribution.

To encourage diverse and non-redundant topics, ECRTM incorporates an embedding clustering regularization term. The overall training loss is defined as

$$\mathcal{L} = \mathcal{L}_{\text{VAE}} + \lambda \cdot \mathcal{L}_{\text{ECR}}, \quad (6)$$

where $\lambda$ is the weight hyperparameter.

#### Supplementary Note 3: Prompt Details for scConcept

**Concept Generation**

```
You are given topics derived from a neural topic model on single-cell RNA-seq.

Each topic includes ONLY:
- A list of the top 100 genes for that topic,
  sorted from highest weight to lowest weight in the model.

Here are all topics (JSON list):
{topics_json_str}

TASK:
You must perform *biological topic distillation* by analyzing ONLY the top-100 gene lists.

Specifically:
1. Identify topics whose top genes indicate highly similar biological pathways or cell
    states.
2. Merge such topics into unified, biologically coherent concepts.
3. Remove topics that are:
   - too vague (gene list does not form a coherent module),
   - too narrow (dominated by a single outlier gene),
   - biologically uninterpretable given the provided gene patterns.
```

````
For each FINAL biological concept, output:
- "name": a concise 2-4 word biological label.
      * Avoid generic labels like "General Regulation".
      * Prefer pathway- or state-specific terms.
- "description": a 10-30 word natural language description
      * Summarize the biological meaning of this concept.
      * Describe the core pathway, cellular program, or functional state.
      * Must be consistent with the concept name and gene list.
      * Do NOT mention topic IDs, gene counts, or modeling details.
- "genes": EXACTLY 100 representative genes.
      * MUST be chosen from the union of gene lists of the merged source topics.
      * MUST reflect a coherent pathway or co-expression program.
      * MUST respect the original sorted importance: higher-ranked genes are preferred.
      * DO NOT invent gene names.
- "source_topics": list of topic IDs merged into this concept.

OUTPUT FORMAT (STRICT):
Respond ONLY with a valid JSON object in EXACTLY this form:

{
  "concepts": [
    {
      "name": "Example Name",
      "description":
        "A concise biological description summarizing the core pathway
        or cellular program represented by this concept.",
      "genes":
        ["GENE1", "GENE2", "GENE3", ..., "GENE100"],
      "source_topics":
        ["topic_0", "topic_3"]
    }
  ]
}

ABSOLUTE RULES:
- DO NOT output anything outside the JSON.
- DO NOT add code fences (such as ```json).
- DO NOT invent genes.
- DO NOT reorder the topics_json_str; use it as provided.
- The final genes must reflect the most informative subset of the top 100 genes per topic.
````

57

**Hierarchical Concept Refinement**

```
You are given ONE biological concept derived from a neural topic model on single-cell RNA-
    seq.

IMPORTANT:
```

58

```
This concept has been determined to be HETEROGENEOUS and contains multiple
distinct biological programs. Your task is to split it into meaningful sub-concepts.

The input concept is a JSON object with:
- "name": concept name
- "genes": EXACTLY 100 genes representing this concept (gene program)

Here is the input concept (JSON):
{concept_json}

TASK:
Split this concept into biologically meaningful sub-concepts.

------------------------------------------------------------
SPLIT REQUIREMENTS (MANDATORY)
------------------------------------------------------------

- Produce EXACTLY 2 to 3 sub-concepts.
- EACH sub-concept must represent a SINGLE, coherent biological program.
- Sub-concepts must be generalizable and interpretable.

For EACH sub-concept:
- "name": concise 2-4 word biological label.
- "description": a 10-30 word natural language description
      * Summarize the biological meaning of this sub-concept.
      * Describe the core pathway, cellular program, or functional state.
      * Must be consistent with the sub-concept name and gene list.
      * Do NOT mention modeling details or the parent concept.
- "genes": select between 10 and 30 genes.
- ALL genes MUST be chosen from the parent concept's gene list.
- DO NOT invent genes.
- DO NOT add genes not present in the parent list.

------------------------------------------------------------
OUTPUT FORMAT (STRICT JSON ONLY)
------------------------------------------------------------

Respond with EXACTLY one JSON object in the following format:

{{
  "concept_name": "<parent_concept_name>",
  "split": true,
  "sub_concepts": [
    {{
      "name": "<sub_concept_name>",
      "description": "A concise description summarizing the biological program represented by
     this sub-concept.",
      "genes": ["GENE1", "GENE2", ...]
```

```
    }}
  ]
}}

OUTPUT RULES:
- "split" MUST be true.
- "sub_concepts" MUST contain 2-3 items.
- Each "genes" list length MUST be between 10 and 30 (inclusive).
- Every gene MUST appear in the parent concept gene list.
- Respond ONLY with valid JSON.
- Do NOT include explanations, comments, or markdown.
```

60

##### Developmental Concept Extraction

```
You are given topics derived from a neural topic model on single-cell RNA-seq data.

Each topic includes ONLY:
- A list of the top 100 genes for that topic,
  sorted from highest weight to lowest weight in the model.

Here are all topics (JSON list):
{topics_json_str}

------------------------------------------------------------
TASK: Developmental Potency Program Distillation
------------------------------------------------------------

Your goal is to identify biologically coherent gene programs that reflect
DIFFERENT LEVELS OF CELLULAR DEVELOPMENTAL POTENCY.

You must analyze ONLY the provided top-100 gene lists from the topics.
No external knowledge, marker lists, or invented genes are allowed.

The target developmental potency categories are EXACTLY the following six:

1. Toti.  (totipotent)
2. Pluri. (pluripotent)
3. Multi. (multipotent)
4. Oligo. (oligopotent)
5. Uni.   (unipotent)
6. Diff.  (differentiated)

IMPORTANT:
- These categories define an INTERPRETIVE FRAMEWORK, not guaranteed labels.
- Some categories MAY NOT be detectable in the provided topics.
- If no coherent gene program exists for a category, you MUST output an empty gene list for
   that category.
```

61

```
-----------------------------------------------------------
BIOLOGICAL INTERPRETATION GUIDELINES
-----------------------------------------------------------

You should infer developmental potency ONLY from gene patterns such as:
- stemness vs lineage restriction,
- transcriptional plasticity vs functional specialization,
- progenitor-like vs terminal differentiation programs,
- broad developmental regulators vs lineage-specific effector genes.

DO NOT:
- assume cell types,
- assume developmental time points,
- use known CytoTRACE markers explicitly,
- force topics into categories if evidence is weak.

A gene program must show CONSISTENT biological signals across one or more topics
to be assigned to a potency category.

-----------------------------------------------------------
PROGRAM CONSTRUCTION RULES
-----------------------------------------------------------

For EACH of the six potency categories:

- Identify zero or more topics whose top genes SUPPORT that potency level.
- Extract a representative gene set from those topics.

Gene selection rules:
- Genes MUST come ONLY from the union of genes in the selected source topics.
- Prefer genes that are:
  * highly ranked in their topics,
  * recurrent across topics,
  * biologically consistent with the inferred potency level.
- DO NOT invent gene names.
- Gene lists may have VARIABLE length (including empty lists).

-----------------------------------------------------------
OUTPUT FORMAT (STRICT JSON)
-----------------------------------------------------------

Respond ONLY with a valid JSON object in EXACTLY the following format:

{{
  "concepts": [
    {{
      "name": "Toti.",
```

```
      "genes": ["GENE1", "GENE2", "..."],
      "source_topics": ["topic_1", "topic_7"]
    }},
    {{
      "name": "Pluri.",
      "genes": [],
      "source_topics": []
    }},
    {{
      "name": "Multi.",
      "genes": ["GENE_A", "GENE_B"],
      "source_topics": ["topic_3"]
    }},
    {{
      "name": "Oligo.",
      "genes": ["GENE_X", "GENE_Y"],
      "source_topics": ["topic_5", "topic_9"]
    }},
    {{
      "name": "Uni.",
      "genes": ["GENE_M", "GENE_N"],
      "source_topics": ["topic_12"]
    }},
    {{
      "name": "Diff.",
      "genes": ["GENE_D1", "GENE_D2"],
      "source_topics": ["topic_15", "topic_18"]
    }}
  ]
}}

------------------------------------------------------------
ABSOLUTE RULES
------------------------------------------------------------

- DO NOT output anything outside the JSON object.
- DO NOT add code fences (no ```json).
- DO NOT invent genes or modify gene symbols.
- DO NOT force assignments when evidence is weak.
- Empty gene lists are VALID and encouraged when appropriate.
- Interpret potency as a CONTINUUM, not discrete cell types.
```

63

#### Coherence Evaluation

```
You are acting as a senior human expert in single-cell transcriptomics
and systems biology, with extensive experience in interpreting gene modules
derived from topic models.
```

64

```
Your task is to assess the BIOLOGICAL COHERENCE of a gene topic,
as a human expert would do during manuscript review.

Rate the topic's biological coherence on a 1-5 scale:

5 = Very coherent: genes strongly support ONE clear biological program,
    pathway, or cell identity.
4 = Coherent: a clear main program with minor noise.
3 = Moderate: weak or mixed biological signal.
2 = Low: little shared biological meaning.
1 = Not interpretable: essentially random or uninformative genes.

Rules:
- Use ONLY the provided gene list.
- Do NOT invent genes or rely on external unstated data.
- If evidence is insufficient, favor LOWER scores.
- Do NOT provide explanations or comments.

Output STRICT JSON only, in exactly this format:
{{
  "topic_id": "{topic_id}",
  "coherence_score": 3
}}

Topic ID:
{topic_id}

Genes:
{gene_list}
```

### 2 Supplementary Figures

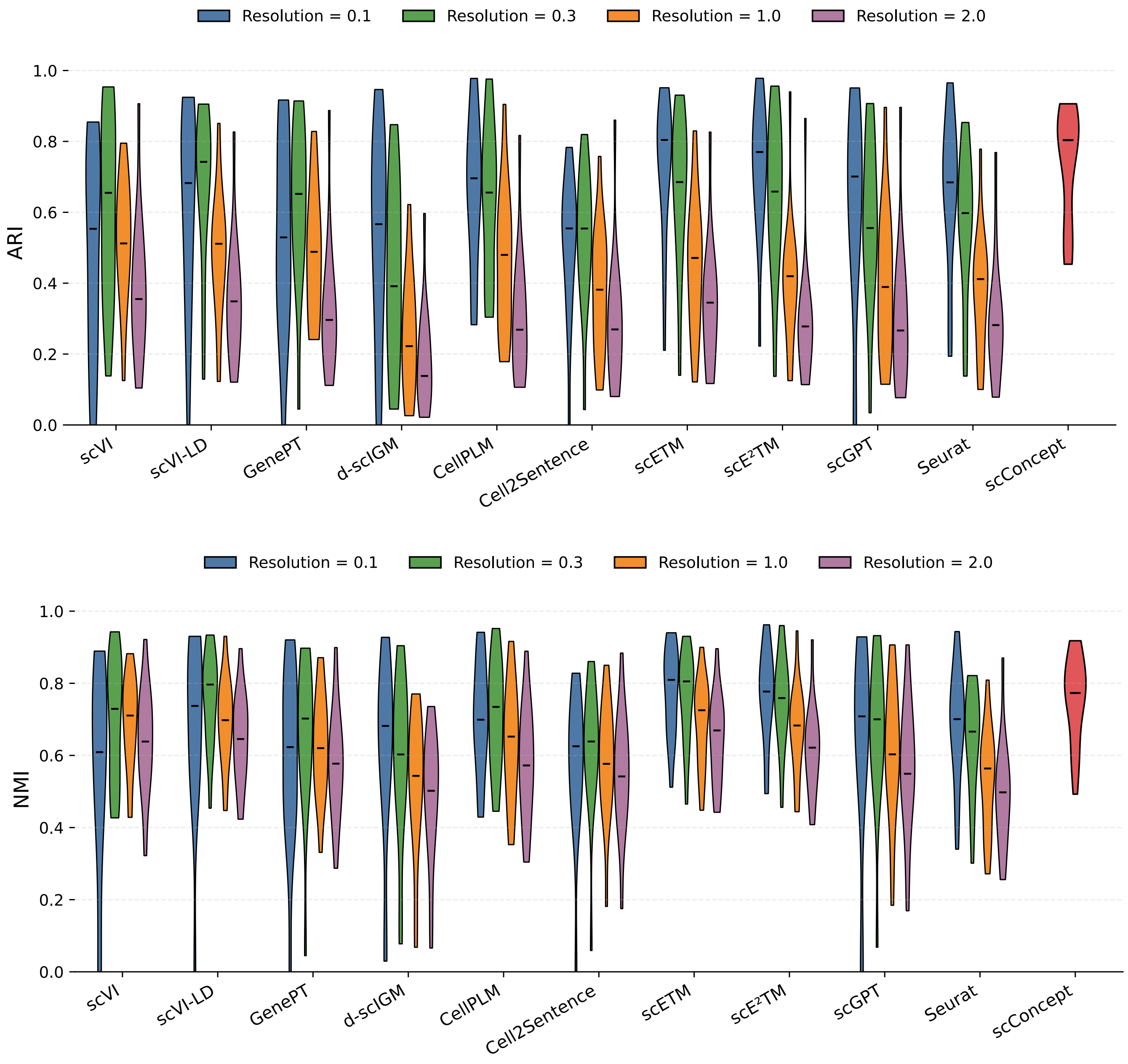

Figure 1: **Clustering performance across different Louvain resolutions.** Clustering accuracy of scConcept and baseline methods evaluated under varying Louvain resolution parameters. For baseline methods, cell embeddings were clustered using the Louvain algorithm with commonly used resolution values (0.1, 0.3, 1.0, and 2.0). Performance was assessed using ARI (top) and NMI (bottom). Each violin plot summarizes the distribution of scores across resolution settings across all datasets.

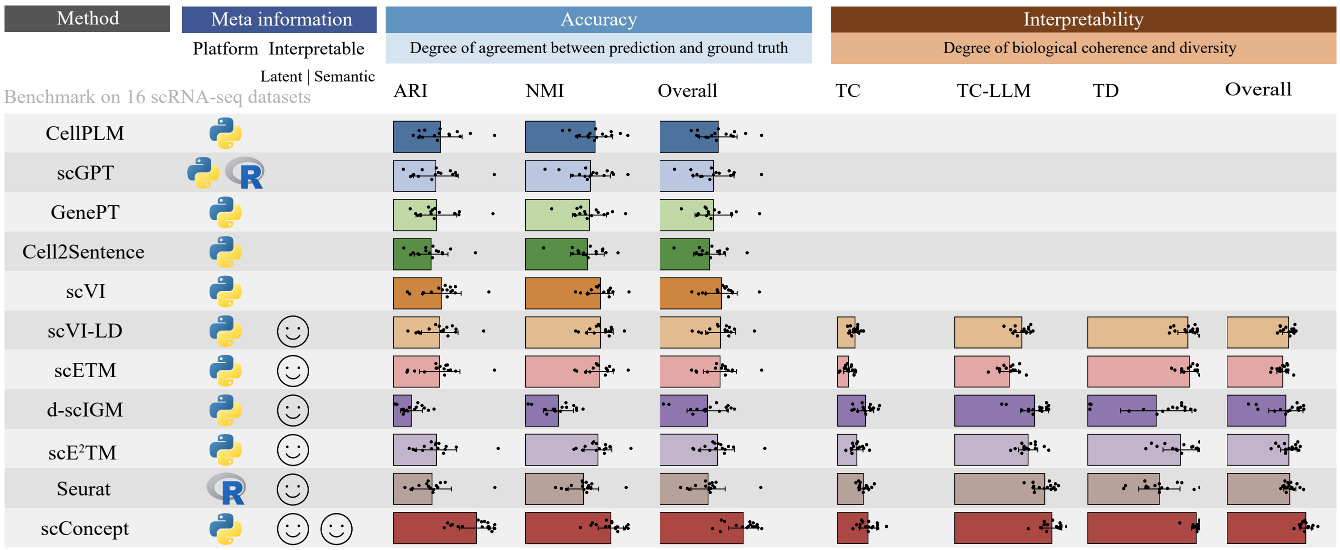

Figure 2: **Performance comparison with reduced embedding dimensionality.** Clustering accuracy and interpretability of scConcept and baseline methods when the embedding dimensionality is set to 10.

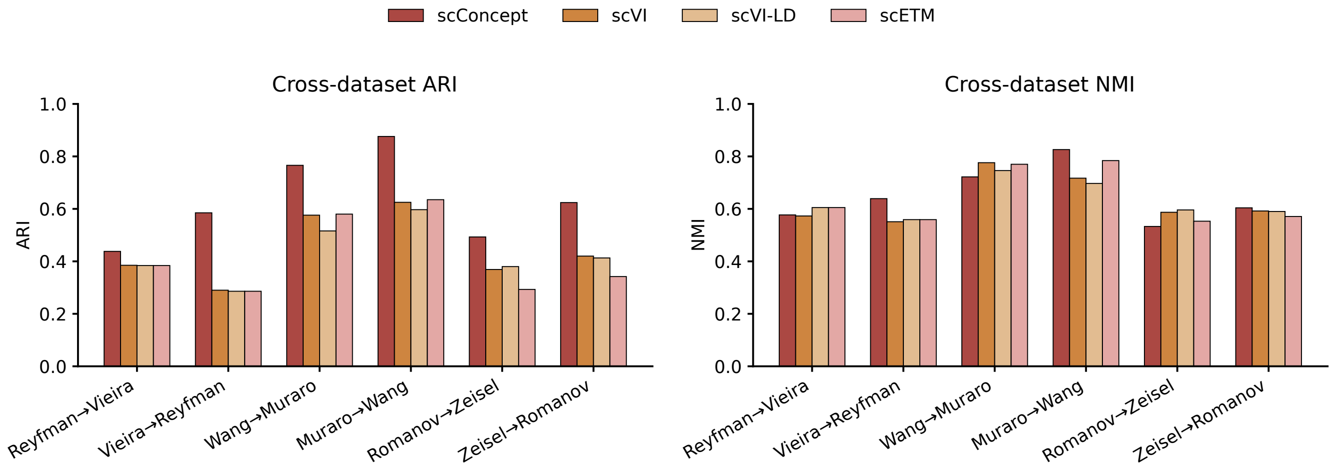

Figure 3: **Cross-dataset clustering performance.** Clustering accuracy of scConcept and baseline methods evaluated across datasets. Models were applied across different datasets to assess generalization under distribution shift. Performance was measured using ARI (left) and NMI (right). Each pair on the x-axis denotes the source and target datasets used for evaluation, demonstrating the robustness of scConcept to cross-dataset variability.

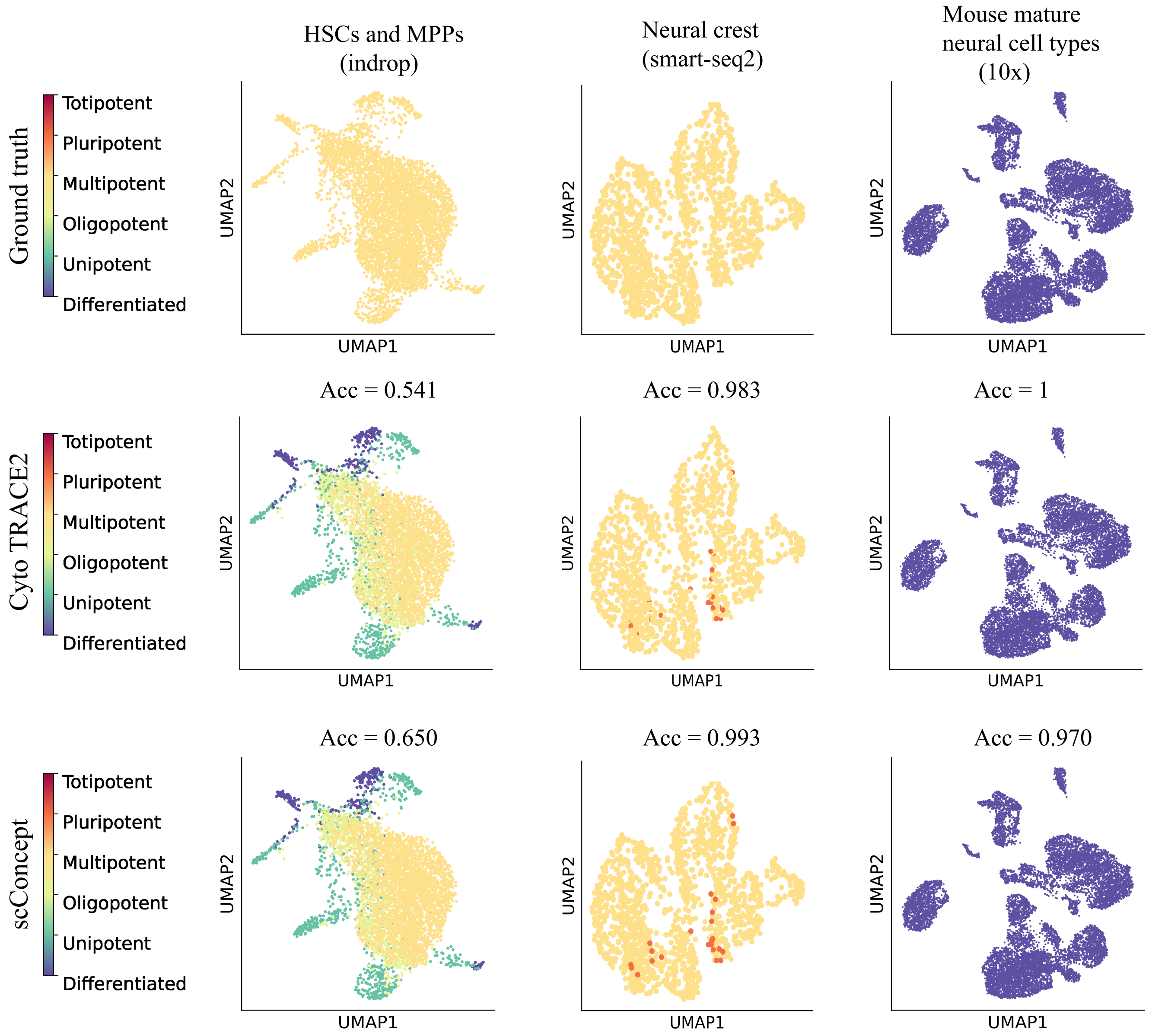

Figure 4: **Developmental potential prediction.** UMAP visualization of three datasets colored by ground-truth developmental annotations (top), CytoTRACE2 predictions (middle), and scConcept predictions (bottom). Prediction performance is evaluated using accuracy (Acc). As each dataset contains only a single dominant developmental stage, rank-based metrics such as Kendall correlation are not applicable.

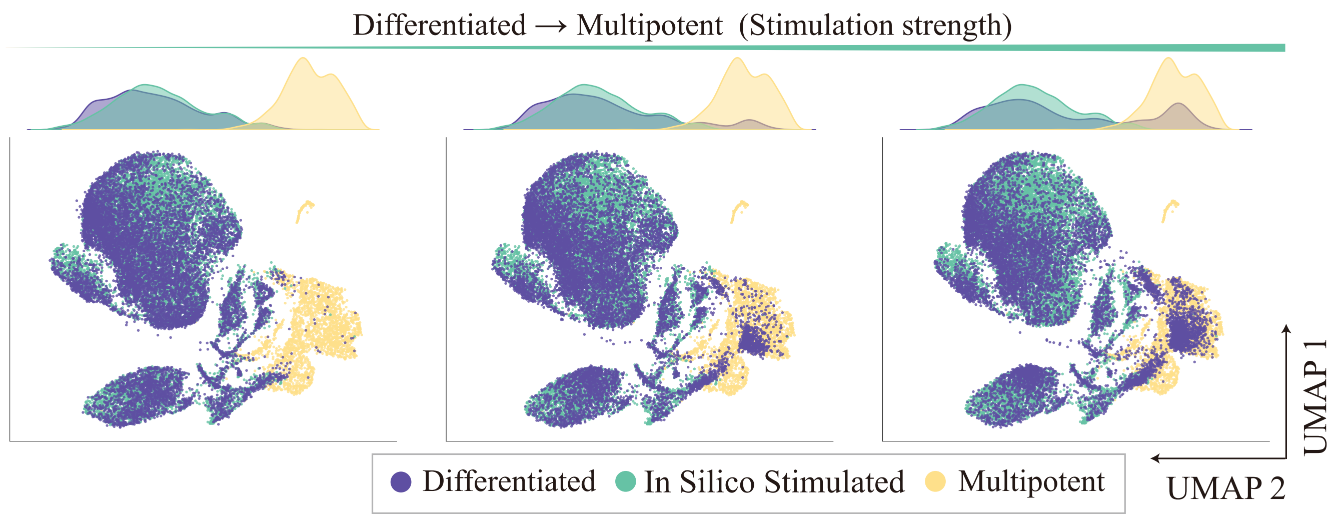

Figure 5: **Concept-driven cell-state transitions.** Cells are perturbed by downregulating genes associated with the *differentiated* concept and upregulating genes associated with the *multipotent* concept, simulating transitions from differentiated to multipotent states. UMAP projections and density plots show progressive shifts in cell states with increasing perturbation strength.

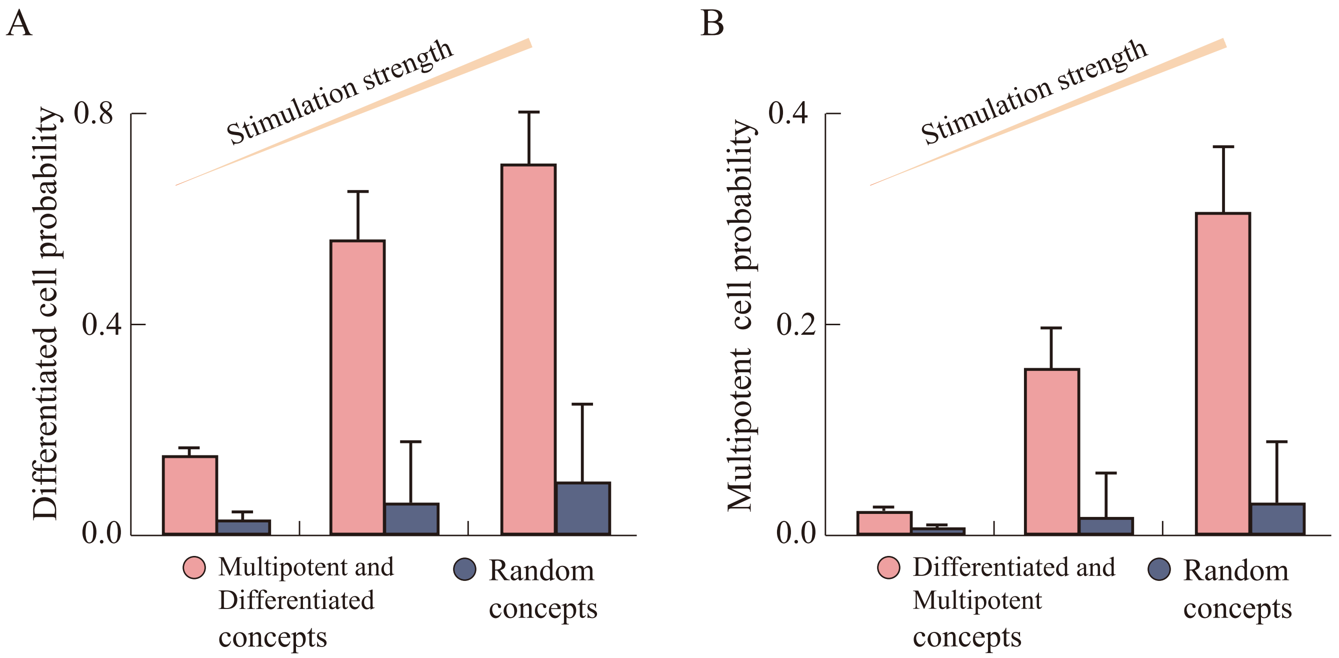

Figure 6: **Classifier-based quantification of concept-driven cell-state transitions.** (A) Probability of cells being classified as differentiated increases as genes associated with the *multipotent* concept are downregulated and genes associated with the *differentiated* concept are upregulated. (B) Probability of cells being classified as multipotent increases under the reverse perturbation, in which genes associated with the *differentiated* concept are downregulated and genes associated with the *multipotent* concept are upregulated. Results are compared with random concept perturbations as a negative control.

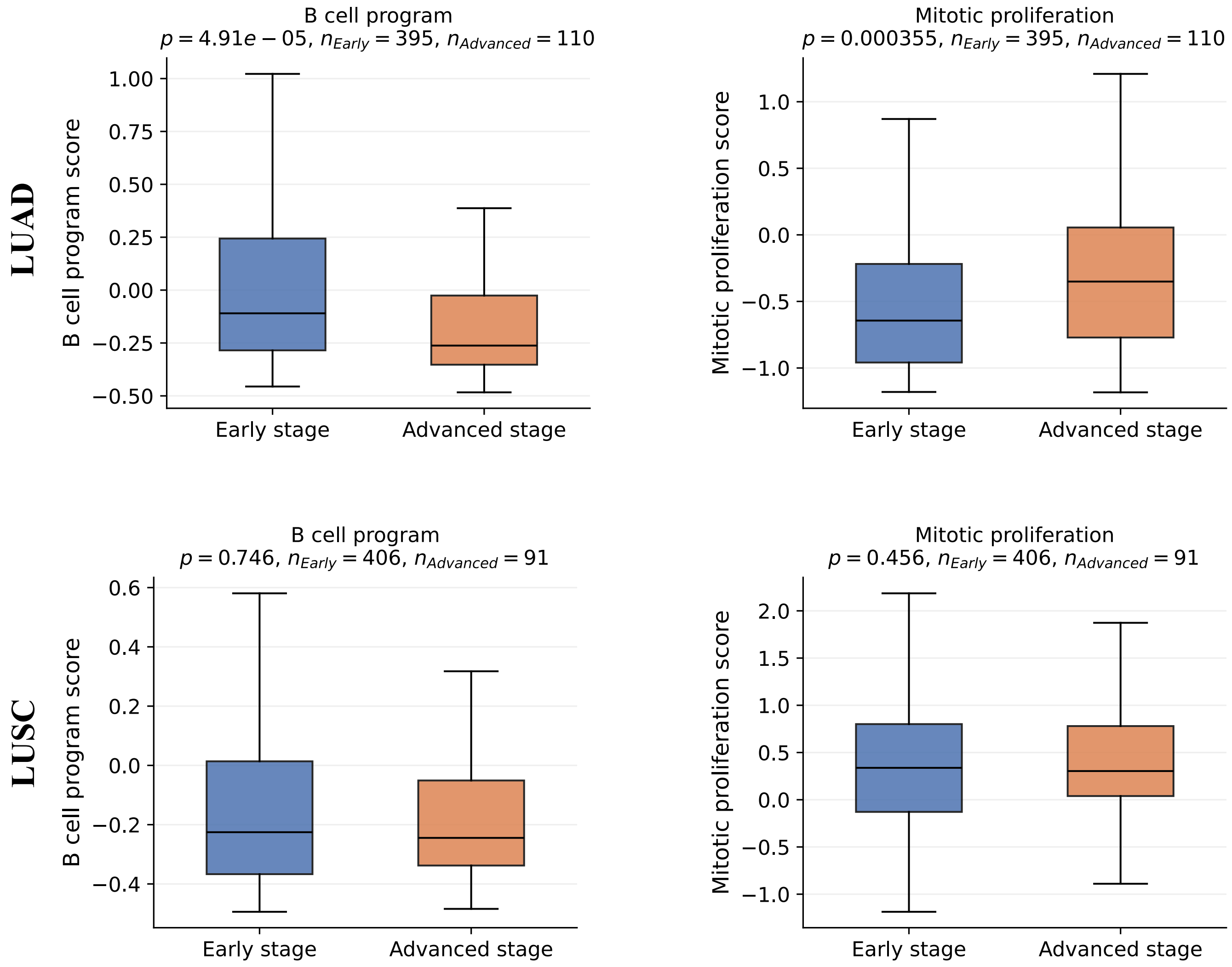

Figure 7: **Association between concept expression and tumor stage.** Expression of the *B cell program* (left) and *Mitotic proliferation* (right) concepts in early- and advanced-stage tumors for LUAD (top) and LUSC (bottom) cohorts. Statistical significance was assessed using the Mann–Whitney U test. Sample sizes for each group are indicated in the panels.

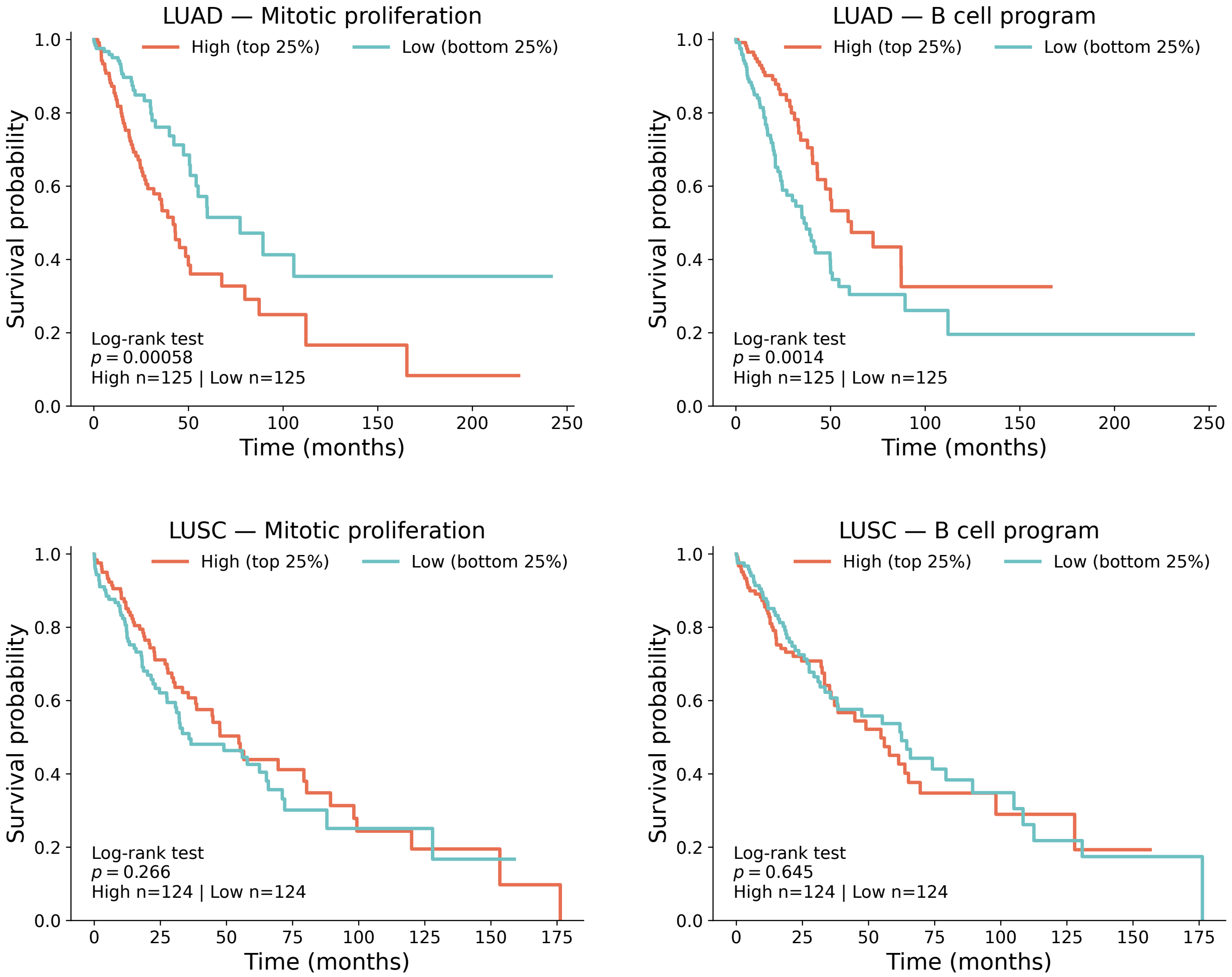

Figure 8: **Association between concept expression and patient survival.** Kaplan–Meier survival analysis of patients stratified by high (top 25%) versus low (bottom 25%) expression of the *Mitotic proliferation* (left) and *B cell program* (right) concepts in LUAD (top) and LUSC (bottom) cohorts. Statistical significance was assessed using the log-rank test. Sample sizes for each group are indicated in the panels.

### 3 Supplementary Tables

Table 1: scRNA-seq datasets with developmental potency annotations.

| Dataset | Accession ID | Protocol | #Sample size | #Broad potency levels | Reference |
|---|---|---|---|---|---|
| Mouse embryo 2 (Smart-seq2) | E-MTAB-3321 | Smart-seq2 | 124 | 2 | Goolam *et al* [3] |
| Mouse mature neural cell types (10x) | GSE249416 | 10x | 11,688 | 1 | Zheng *et al* [4] |
| Neural crest (Smart-seq2) | GSE162044 | Smart-seq2 | 1,804 | 1 | Zalc *et al* [5] |
| HSCs and MPPs (inDrop) | GSE90742 | inDrop | 4,941 | 1 | Rodriguez-Fraticelli *et al* [6] |
| Retinal neurons (10x) | GSE122466 | 10x | 5,347 | 3 | Lo Giudice *et al* [7] |
| Mouse neurogenesis (10x) | GSE249416 | 10x | 14,835 | 3 | Zheng *et al* [4] |

Table 2: Developmental potential concepts identified by scConcept from the Mouse neurogenesis (10x) dataset.

| **Concept name** | **Number of genes** | **Concept genes** |
|---|---|---|
| Totipotent | 0 | – |
| Pluripotent | 0 | – |
| Multipotent | 34 | HES1, HES5, SOX2, SOX3, SOX1, NOTCH1, RFX4, FABP7, LPAR1, PAX6, HOPX, HMGA2, PRDM16, TFAP2C, ZFP36L1, WWTR1, ALDOC, GAS1, ID1, NES, CCND2, CDK6, CLSPN, UHRF1, MCM5, MCM3, BRCA1, CHEK1, DTL, TK1, GMNN, POLE, FGFR3, HELLS |
| Oligopotent | 21 | EOMES, NEUROG2, NEUROD1, SSTR2, UNC5D, EPHA3, ROBO2, DCC, KIF26B, SHISA2, DMRTA2, DACH1, ELAVL4, TCF12, PLXNA4, CELSR1, SLC17A6, NEUROG1, THRB, RFTN2, EYA1 |
| Unipotent | 16 | DLX1, DLX2, DLX5, DLX1AS, DLX6OS1, ARX, LHX6, GAD1, GAD2, SP9, SLC6A1, SLC32A1, NETO1, NKX2-1, POU3F4, TOX2 |
| Differentiated | 44 | LRFN5, LRRTM4, CDH13, OPCML, NRXN1, NLGN1, PTPRD, RYR3, DLG2, RALYL, CADM2, DAB1, EDIL3, PHACTR1, FGF14, TMEM108, KIRREL3, TENM1, TENM2, PCDH9, NAV3, SLC8A1, ANK3, GRIA1, GABRB1, GABRB2, DPP10, CALN1, CACNA1E, CACNA1D, SYBU, RIMBP2, MEF2C, ZFPM2, NRG3, CHL1, MDGA2, HS3ST4, PCDH7, EPHA5, NXPH3, NECAB1, PDZD2, ADGRB3 |

Table 3: Literature support for the top 10 genes in the *Mitotic proliferation* concept identified by scConcept in lung cancer. Most genes have prior evidence linking them to lung cancer progression or prognosis.

| **Index** | **Gene name** | **Gene stable ID** | **Reference** |
|---|---|---|---|
| 1 | PIMREG | ENSG00000129195 | Jiang *et al* [8] |
| 2 | DEPDC1 | ENSG00000024526 | Wang *et al* [9] |
| 3 | KIF20A | ENSG00000112984 | Wang *et al* [10] |
| 4 | HJURP | ENSG00000123485 | Tan *et al* [11] |
| 5 | SPC25 | ENSG00000152253 | Chen *et al* [12] |
| 6 | AURKB | ENSG00000178999 | Bertran-Alamillo *et al* [13] |
| 7 | NEK2 | ENSG00000117650 | Shi *et al* [14] |
| 8 | H3C8 | ENSG00000273983 | Not reported |
| 9 | GTSE1 | ENSG00000075218 | Zhang *et al* [15] |
| 10 | PBK | ENSG00000168078 | Shih *et al* [16] |

2. Hanna Wallach, David Mimno, and Andrew McCallum. Rethinking lda: Why priors matter. *Advances in neural information processing systems*, 22, 2009.
